# Novel insights into hippocampal perfusion using high-resolution, multi-modal 7T MRI

**DOI:** 10.1101/2023.07.19.549533

**Authors:** Roy A. M. Haast, Sriranga Kashyap, Dimo Ivanov, Mohamed D. Yousif, Jordan DeKraker, Benedikt A. Poser, Ali R. Khan

## Abstract

We present a comprehensive study on the non-invasive measurement of hippocampal perfusion. Using high-resolution 7 Tesla arterial spin labelling data, we generated robust perfusion maps and observed significant variations in perfusion among hippocampal subfields, with CA1 exhibiting the lowest perfusion levels. Notably, these perfusion differences were robust and detectable even within five minutes and just fifty perfusion-weighted images per subject. To understand the underlying factors, we examined the influence of image quality metrics, various tissue microstructure and morphometry properties, macrovasculature and cytoarchitecture. We observed higher perfusion in regions located closer to arteries, demonstrating the influence of vascular proximity on hippocampal perfusion. Moreover, *ex vivo* cytoarchitectonic features based on neuronal density differences appeared to correlate stronger with hippocampal perfusion than morphometric measures like gray matter thickness. These findings emphasize the interplay between microvasculature, macrovasculature, and metabolic demand in shaping hippocampal perfusion. Our study expands the current understanding of hippocampal physiology and its relevance to neurological disorders. By providing *in vivo* evidence of perfusion differences between hippocampal subfields, our findings have implications for diagnosis and potential therapeutic interventions. In conclusion, our study provides a valuable resource for extensively characterising hippocampal perfusion.

## Introduction

The brain’s multi-scale organisation enables processing of different sensory inputs through pathways optimised for storing, updating, and recollecting relevant information^1^. In particular, the structure and function of the hippocampus (or ‘hippocampal formation’) have been at the centre of attention in a plethora of studies focused on the brain and cognitive aging, especially those investigating memory (dys)function, where it was found to be involved in episodic memory (i.e., encoding and retrieval of information tied to a specific time and place), as well as in other types of declarative memory^2^.

Although the hippocampus has been studied as a singular region for several years, emerging *in vivo* imaging (e.g., ultra-high field MRI) and analysis (e.g., topologically-correct unfolding) methods have enabled a better appreciation of its internal organisation^3,4^. While a lot is known about the histological sub-divisions of the hippocampal formation^5^, several studies have provided *in vivo* evidence of hippocampal subfields namely, the subiculum (Sub), the Cornu Ammonis (CA) fields 1-4 and Dentate Gyrus (DG), their unique anatomical properties^6,7^ and their distinct roles in memory processing^8,9^ and sensitivity to age-related changes^10–13^. The fact that there are subfield-specific properties likely render differential effects across diseases^14^ and disease subtypes such as those observed in focal epilepsy^15^. Unfortunately, the neurobiological substrates underlying age- or disease-related changes across and between subfields remain poorly understood.

Hippocampal anatomy varies considerably between individuals^16^ and its fine details are indistinguishable using standard anatomical T_1_-weighted scans. In most cases, specialised coronal T_2_-weighted scans with a high in-plane resolution positioned oblique to the hippocampus’s long axis are used to delineate its convoluted anatomy^17^. Regardless, out-of-plane issues during manual, voxel-based labelling procedures renders it difficult to respect topological constraints such as the contiguity of hippocampal subfields^4,18^. Looking closer into its structure, there are spatial differences in cytoarchitecture^3^ and vascular density^19^. Taking together, the patterns of vascularisation and perfusion across the hippocampal tissue suggests the selective vulnerability of hippocampal regions for vascular pathologies^14,20^ and implicate its role in lifelong exposure to risk factors on hippocampal integrity^21^.

Recent efforts using high-resolution time-of-flight magnetic resonance angiography (TOF-MRA) data enabled differentiation of hippocampal vascularisation patterns and assessment of their impact on cognitive functioning in cerebral small vessel disease patients^22,23^. Similarly, ferumoxytol-enhanced susceptibility-weighted imaging revealed differences across subfields in terms of microvascular density^24^. Nonetheless, it has remained unclear how these macro- and microvascularisation patterns translate to variability in the amount of blood (in ml/100g/min) perfused in the hippocampal tissue. Inherent to the challenges in characterisation of the hippocampal structure, measuring perfusion non-invasively (without contrast agents) *in vivo* for detailed quantification of human hippocampal perfusion has been thus far unexplored.

Arterial spin labelling (ASL) is a non-invasive MRI method that allows quantitative measurements of cerebral perfusion^25^. ASL relies on arterial blood water as endogenous tracer but typically suffers from low signal-to-noise ratio (SNR) due to low grey matter microvascular density relative to the tissue volume^26^. At 3 Tesla (3T), averaging tens of images acquired in roughly 2-4 minutes can provide a low-resolution (4 mm isotropic) perfusion map using ASL. However, this is insufficient to delineate perfusion differences across the hippocampus which has a more fine-grained neuroanatomical composition. This fine-grained anatomical structure in addition to the inter-subject variability^16^ limits the ability to perform simple across-subjects averaging to improve SNR of the data.

In this study, we tackle the aforementioned challenges in characterising hippocampal perfusion by capitalising upon advances in acquisition and analysis strategies. For our first goal, we optimised an ASL acquisition scheme at 7T and leverage the gain in SNR with field strength and lengthening T_1_ at UHF to obtain robust high-resolution (1.5 mm^3^) hippocampal perfusion data^27,28^. The characterisation of the hippocampal anatomy is facilitated using sub-millimetre resolution T_2_-weighted data and construction of a surface-based representation using a novel approach, HippUnfold^29^. By modelling the hippocampus as a folded surface, this approach circumvents issues experienced with manual voxel-based methods and enables inter-subject alignment, as well as parcellation based on a subfield atlas that respect topological constraints^4^. Joint application of these methods enable a spatially-precise characterisation of tissue perfusion across its grey matter and subfields, in particular. For our second goal, we assess the impact of nearby arterial structures reconstructed using a high-resolution TOF-MRA on the perfusion maps. And finally, by leveraging the harmonized unfolded space, we will assess the cross-correlation between hippocampal perfusion, morphometry, other MRI-based properties as well as cytoarchitectonic features extracted from a histological hippocampal sample provided by the BigBrain project^30,31^. Altogether, the presented results provide novel, significant neuroscientific findings that will aid the community to interpret hippocampal (subfield) changes relevant in the context of neurological diseases and/or cognitive neuroscience, as well as an imaging framework that can be used to guide researchers in setting up protocols and analysis of such data to study hippocampal perfusion.

## Results

### Perfusion in the hippocampus

Subject-specific quantitative perfusion data were mapped onto their hippocampal mid-thickness surfaces reconstructed using the HippUnfold output for its in-depth characterisation. Surface mapping and unfolding demonstrate that the hippocampal grey matter is characterised by a spatially varying perfusion pattern. The average perfusion map across all 8 runs, 10 subjects and both hemispheres (i.e., totalling to an average of 160 perfusion maps) depicted in Fig. 1A, showcased the following patterns: (a) lower perfusion in the anterior portion (hippocampal head) and along the hippocampal sulcus (white arrows), (b) higher perfusion towards the posterior portion (hippocampal tail) and both proximally and distally (solid arrow) from its boundary with the neocortical tissue.

**Figure 1.**
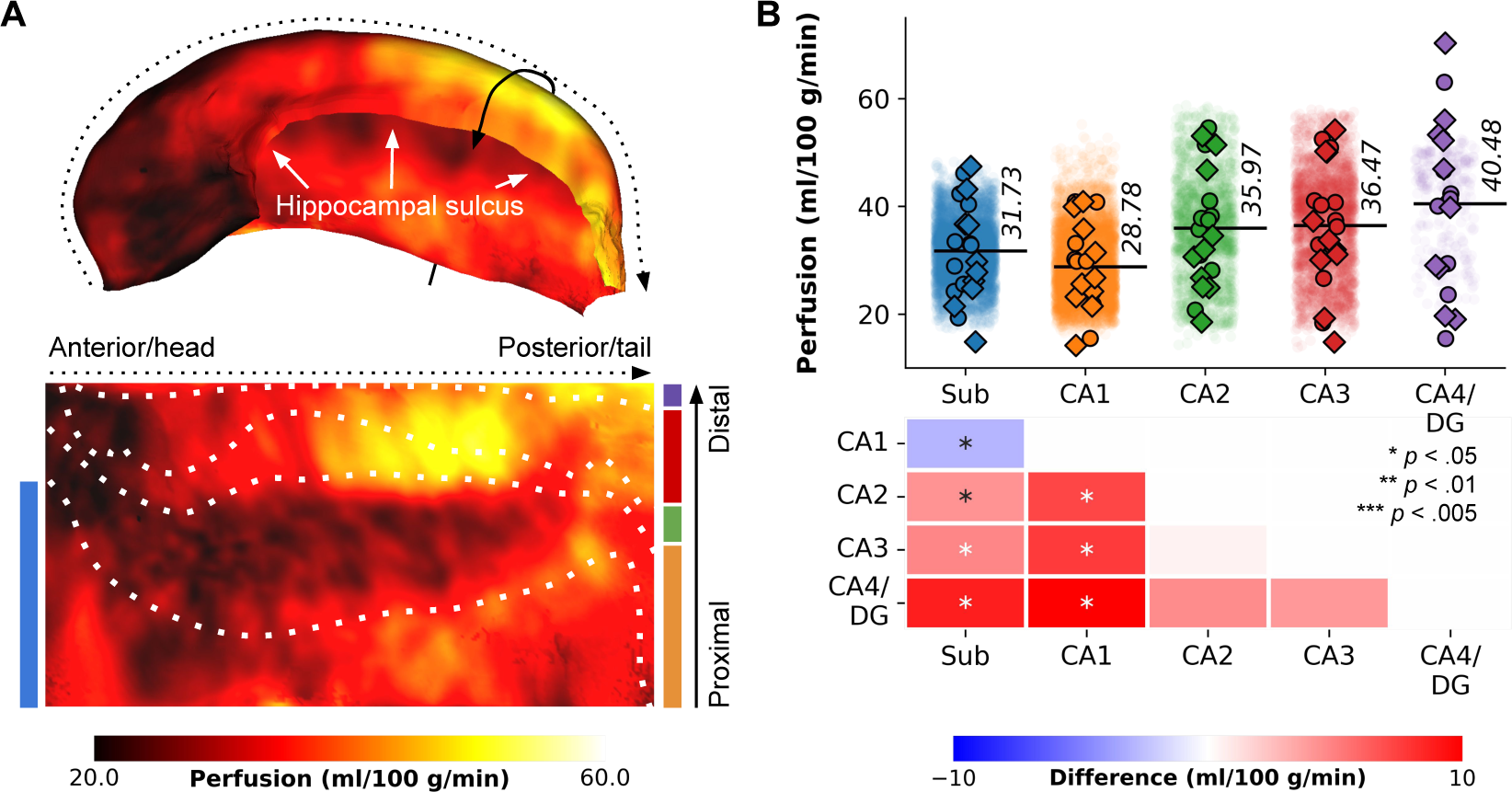
Perfusion mapping in the hippocampus. (A) The figure displays perfusion values (ml/100 g/min) mapped on folded and unfolded hippocampal surfaces. The dotted and solid arrows indicate the anterior-posterior and proximal-distal axes, respectively. Subfield boundaries, derived from cytoarchitectonic features of the BigBrain atlas, are overlaid on the unfolded map. (B) Subfield averages, color-coded based on the subfield atlas overlaid on maps in D, are presented for each subject and hemisphere (circles for the left hemisphere, diamonds for the right hemisphere), as well as per vertex (semi-transparent dots averaged across subjects and hemispheres). Pairwise comparisons between subfield averages are depicted as heatmaps, with FDR*_BH_*-corrected p-values indicated by asterisks: *p < .05, **p < .01, ***p < .005.

To facilitate interpretation, the perfusion map was mapped onto the canonical unfolded hippocampal surface, with the Sub, CA1, CA2, CA3 and CA4/DG subfields arranged from bottom to top, and the head, body and tail aligned from left to right. Friedman’s tests for repeated measures, based on subject-wise subfield averages, demonstrated significant differences in perfusion values among subfields (*χ*^2^ (4) = 19.68, *p* < .001, Fig. 1B). Particularly, CA1 exhibited significantly lower perfusion (average of 28.78 ml/100 g/min) compared to the other subfields (*p_FDR_ <* .05), as indicated by pairwise comparisons (refer to heat maps in Fig. 1B). Perfusion levels in CA2, CA3, and CA4/DG did not exhibit significant variations among each other. These findings broadly align with Duvernoy’s seminal work^19^. For interactive exploration of vertex- and subfield-wise data, we direct the reader to our online app^1^.

#### Reliability of hippocampal perfusion estimates

The low microvascular density poses challenges for ASL-based perfusion data^26^, resulting in relatively low signal-to-noise ratio (SNR), especially at the required spatial resolutions for hippocampal subfield imaging. To study stability and sensitivity in detecting intra-hippocampal differences, we constructed hippocampal perfusion maps by aggregating data from multiple runs and subjects. Fig. 2 illustrates the evolution of the perfusion map in the hippocampus, providing insights into the minimum required number of runs and/or subjects for future studies. We evaluated the stability of average perfusion and the variability, assessing the coefficient of variation for perfusion values (i.e., homogeneity), across vertices in each subfield, with varying subject (S1-10, Fig. 2A) and run (R1-8, Fig. 2B) quantities (in consecutive order). Fig. 2C presents the unfolded representation of vertex-wise perfusion estimates as a function of included runs (columns) and subjects (rows), with thick outlines and gray background indicating significant subfield effects (*p* < .05 based on the Friedman’s test). Notably, this visual representation demonstrates a gradual transition towards the final perfusion pattern, with discernible subfield effects observed from six subjects onwards. Two key observations emerge from this analysis: (a) the stability of the mean perfusion signal (solid lines) is largely influenced by the number of subjects in the cohort, and (b) perfusion variability (dotted lines) tends to deviate more than the mean perfusion signal, particularly with smaller amounts of included data.

**Figure 2.**
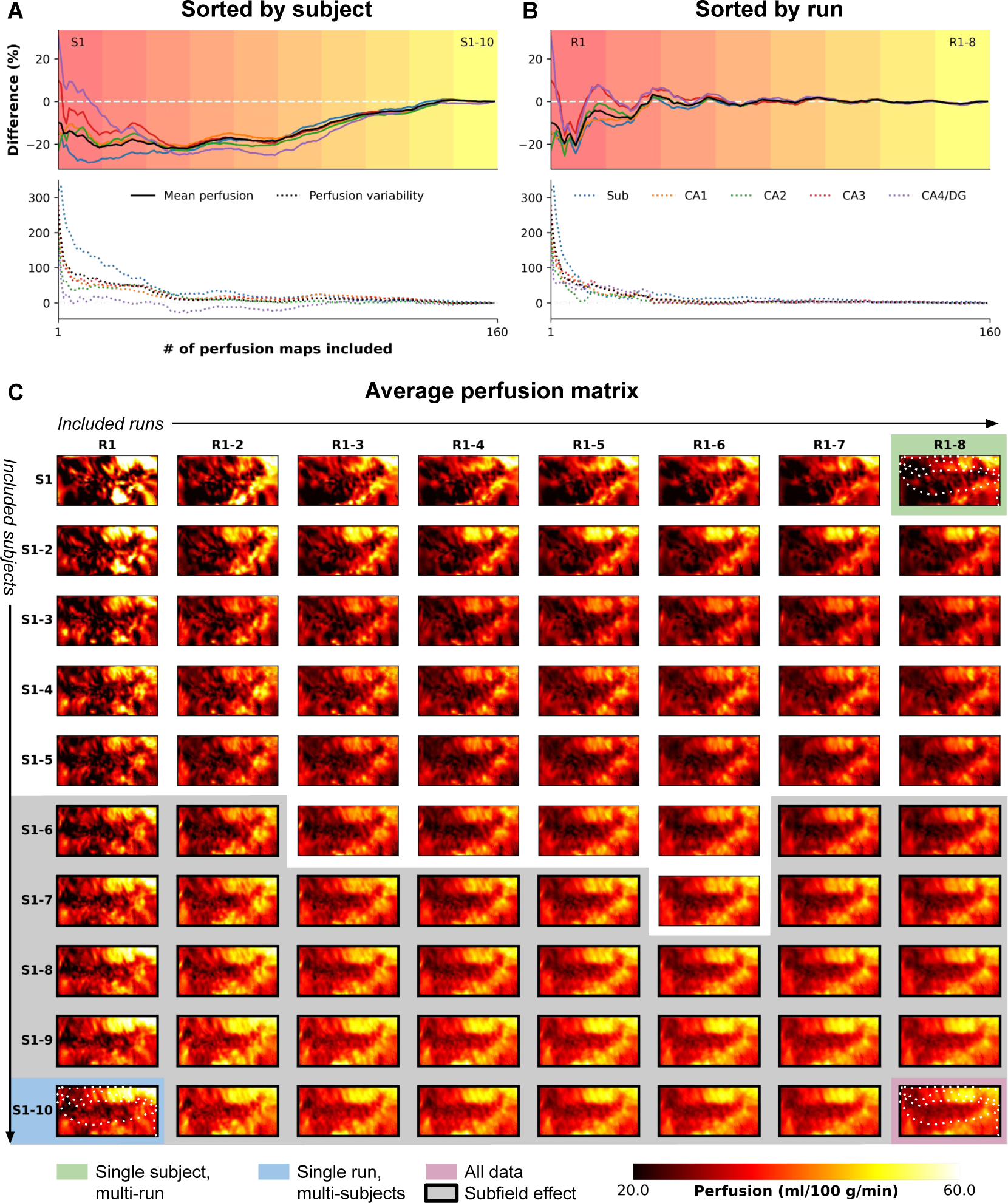
Evolution of high-resolution perfusion maps. The figure illustrates the progression of high-resolution perfusion maps, showcasing the percentage difference in mean perfusion (solid lines, top) and variability (dotted lines, bottom) across the entire hippocampus (in black) and individual subfields (color-coded). The evolution is presented as a function of (A) the number of included subjects or (B) runs. (C) Additionally, the figure depicts the unfolding of the average perfusion map and its evolution as a function of the number of included runs (rows) and subjects (columns) in consecutive order.

The findings in Fig. 2A and B emphasize the critical role of cohort size and data inclusion in achieving stable and reliable hippocampal perfusion measurements, shedding light on the interplay between stability, variability, and the quantity of included data. Furthermore, they reveal variations in the evolution of perfusion estimates depending on the sorting criteria, either subject-wise or run-wise. Therefore, we conducted additional analyses by performing N=1000 iterations, shuffling the order of subjects and runs each time, and calculating the median value to address potential sampling biases (Fig. S1). It is important to note that the left and right hemisphere data were averaged for each iteration, similar to Fig. 2C, as they were acquired simultaneously during a single run. Alongside the mean and variability of perfusion estimates, we assessed the dependence of perfusion temporal SNR (tSNR) and the effect size of between-subfield differences using the Friedman’s test Q-statistic. Heat maps in Fig. S1 show the median across iterations relative to the results obtained from the full fit analysis based on the 160 perfusion maps, indicated by the black round marker in A, which corresponds to the maps presented in Fig. 1. The results confirm a lower dependency of mean perfusion estimates on the amount of included data (up to 1% difference, Fig. S1A) compared to between-vertex variability (up to 100%, Fig. S1B). Notably, the number of included subjects exerts the strongest impact. In contrast, perfusion tSNR exhibits a gradual stabilization with increasing runs, rather than subjects (Fig. S1C). Consistent with improved subfield homogeneity (Fig. S1B), the between-subfield effect size gradually increases as more subjects are included (Fig. S1D), with a significant effect (*p* < .05) already observable starting from three subjects.

### Characterisation of MRI quality and morphometric hippocampal features

To assess the influence of other hippocampal properties on the perfusion pattern, we extracted several acquisition- and morphology-related metrics. The acquisition-related metrics encompassed perfusion time-course stability (tSNR), B^+^ for blood labelling efficiency, tissue T_1_, susceptibility-induced image distortions, and partial volume estimates (PVE) for gray matter (GM), white matter (WM), and cerebrospinal fluid (CSF) tissue classes. The average tSNR of hippocampal perfusion was 3.35 *±* 0.84 (Fig. S2A). The labelling efficiency, represented by B^+^, exhibited an average of 12.03 *±* 1.15 *µ*T across all data points (Fig. S2B) (6.54 *µ*T is required to meet the adiabatic condition for inversion). Susceptibility-induced distortions in perfusion imaging were most prominent in the anterior portion (hippocampal head), peaking at 0.3 mm within the Sub and CA1 (Fig. S2C), consistent with their proximity to the air-tissue interface. However, the magnitude of these distortions was relatively modest compared to those typically observed in functional or diffusion MRI^32,33^.

For validation purposes, we also quantified additional hippocampal tissue properties, including morphology-related metrics such as cortical thickness, curvature, and gyrification derived using HippUnfold, as well as metrics related to underlying tissue microstructure, predominantly reflecting myelin content based on T_1_w/T_2_w ratio maps. Each metric displayed distinct spatial patterns (Fig. S3). Cortical thickness was lowest in CA2, while gyrification was most pronounced along CA1 towards the head. The subiculum exhibited the strongest myelination, as indicated by low T_1_ values but high T_1_w/T_2_w ratios. These findings align with previous observations, confirming the expected variations in hippocampal tissue properties across subfields^29^.

### Tracing hippocampal vascularisation

In our second objective, we utilized high-resolution TOF-MRA data to reconstruct the macrovasculature of the hippocampus for combined analysis with the perfusion data. Fig. 3A illustrates a 3D reconstruction example of the right hippocampal macrovasculature for a single subject. The topology of the reconstructed vasculature aligns closely with previously identified patterns and trees of hippocampal vascularisation^23^. The prominent internal carotid artery (ICA, depicted by magenta solid lines) serves as the primary blood supply source to the medial temporal lobe. The posterior communicating artery (PCA) connects the ICA with the P1 (cyan) and P2 (white) segments of the posterior cerebral artery, with the latter running parallel to the anterior-posterior axis of the hippocampus. Similar to the PCA, the anterior choroidal artery (orange) arises from the ICA and follows a superior position in the same direction. Interactive visualizations of these 3D reconstructions, as well as additional representations such as node-wise networks and vessel geometry properties (including B^+^), specific to each subject’s hippocampus, can be accessed in the interactive HTML notebooks provided in the online code repository^2^. Collectively, these visualizations demonstrate that the network of interconnected arteries described above was identifiable in most cases. However, the detection of thinner arteries, such as the anterior and posterior hippocampal arteries, was less reliable across subjects due to their diameter falling below the effective resolution of the TOF-MRA data.

**Figure 3.**
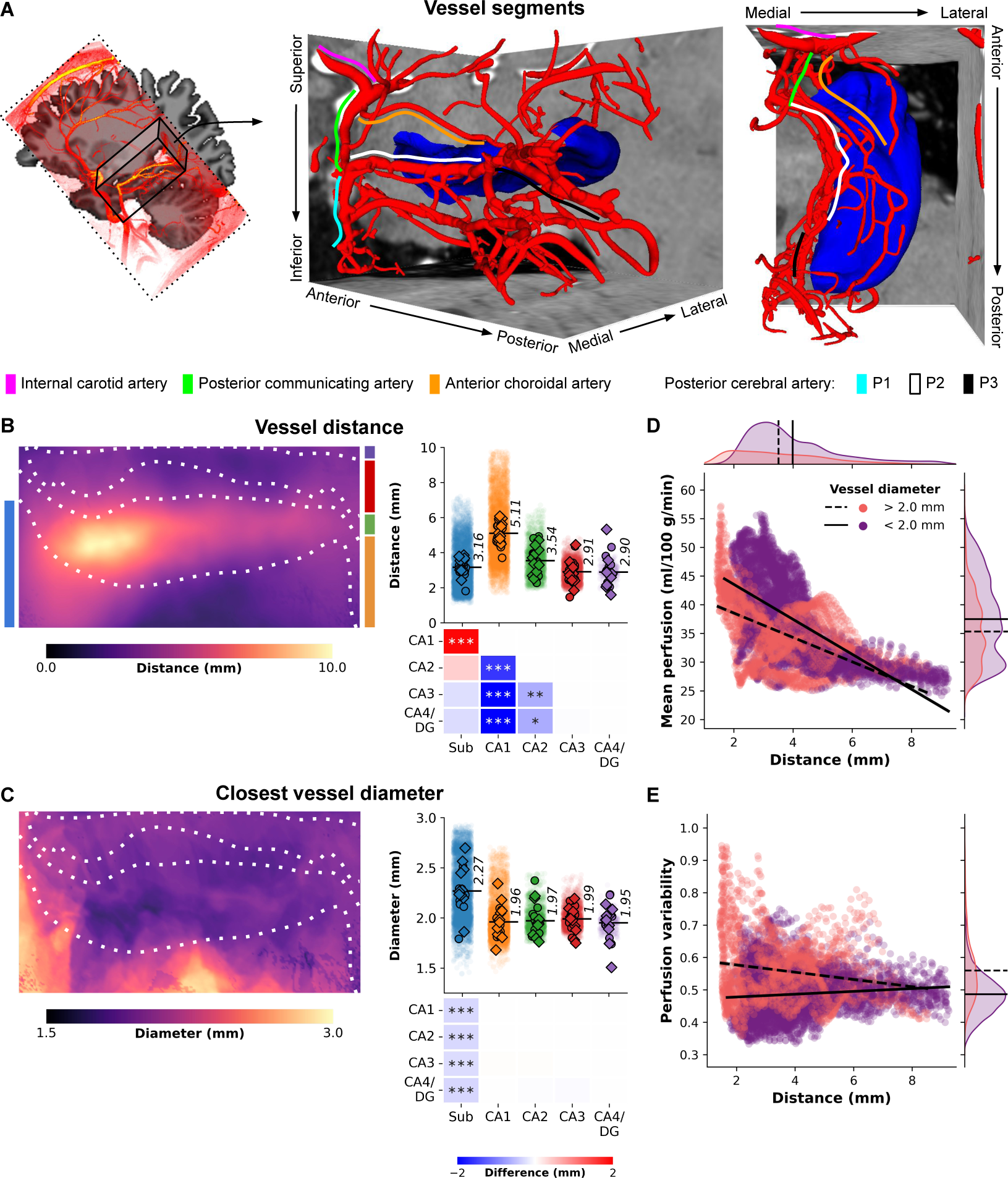
Hippocampal vasculature and perfusion relationship. (A) Three-dimensional reconstruction of a subject’s macrovasculature in close proximity to the right hippocampus, showcasing delineated vessel segments. (B) Hippocampal vessel distance (mm) depicted on an unfolded hippocampal surface. Strip plots display color-coded subfield averages for each subject, including left hemisphere (circles) and right hemisphere (diamonds), along with per vertex values (i.e., averages across subjects and hemispheres shown as semi-transparent dots). Heatmaps illustrate pairwise comparisons between subfield averages, with FDR*_BH_*-corrected p-values indicated by asterisks: *p < .05, **p < .01, ***p < .005. (C) Similar to (B), but representing vessel diameter (mm) of the nearest vessel. (D) Scatter plot illustrating the relationship between vertex-wise mean perfusion (ml/100 g/min) and the shortest distance to a vessel (mm), stratified by respective vessel diameter (color-coded as thinner or thicker than 2 mm). Linear fits for each group are depicted by solid and dashed black lines. (E) Similar to (D), but contrasting with perfusion variability determined by the coefficient of variation across all maps (i.e., across runs and subjects).

#### Linking hippocampal vascularisation and perfusion

Once the vessel tree for each subject was established (Fig. 3A), vessel-related metrics were projected onto the hippocampal surfaces to examine the positioning of vertices and subfields relative to the hippocampal vasculature (refer to Fig. S4A for an example of the metrics). It is important to note that the presented diameter values are estimates limited by the spatial resolution of the TOF-MRA data, which hinders the reliable identification of vessels smaller than 0.5 mm. The averaged results across subjects highlight variations in the distance to vessels throughout the hippocampus, ranging from 0 to 10 mm, with differences observed between subfields (*χ*^2^ (4) = 25.2, *p* < .001, Fig. 3B). Specifically, CA1 is located farthest from the vessels, with an average distance of 5.11 mm, while the subiculum (3.16 mm), CA3 (2.91 mm), and CA4/DG (2.90 mm) exhibit closer proximity to macrovasculature structures. Notably, the vessel distance map (Fig. 3B) demonstrates a distinct pattern along the anterior-posterior axis of the hippocampus. Moreover, vessels in close proximity to the subiculum (e.g., PCA) tend to have relatively larger diameters (average of 2.27 mm) compared to vessels near other subfields (*χ*^2^ (4) = 21.4, *p* < .001, Fig. 3C). An important finding from this analysis is that the largest perfusion values are associated with the proximal vasculature (Fig. 3D), and overall perfusion signals remain relatively stable across different vessel sizes and their distances from hippocampal tissue (Fig. 3E).

### Quantification of hippocampal features cross-correlation

Having demonstrated that perfusion levels vary across hippocampal grey matter, with higher perfusion levels linked to a closer distance to vascular structures, and that we have confirmed previously established patterns for its morphometry (thickness, gyrification and curvature) and myelination^34^, we set out to map out their interdependencies. To accomplish this, we computed the Pearson’s correlation coefficient to assess the similarity among all pairs of hippocampal features and their vertex-wise averages (see Fig. 4A). Perfusion did not demonstrate significant correlations with image distortion, B^+^, T_1_, and partial volume estimates (*p_roll_ >* .05, Fig. 4A). However, a significant correlation was observed with tSNR (Pearson’s *r* = .81, *p_roll_ <* .001, Fig. 4A), indicating that higher perfusion values were obtained in regions less dominated by noise. It is reassuring that our measurements and thus findings of hippocampal perfusion patterns appear robust to acquisition-related metrics.

**Figure 4.**
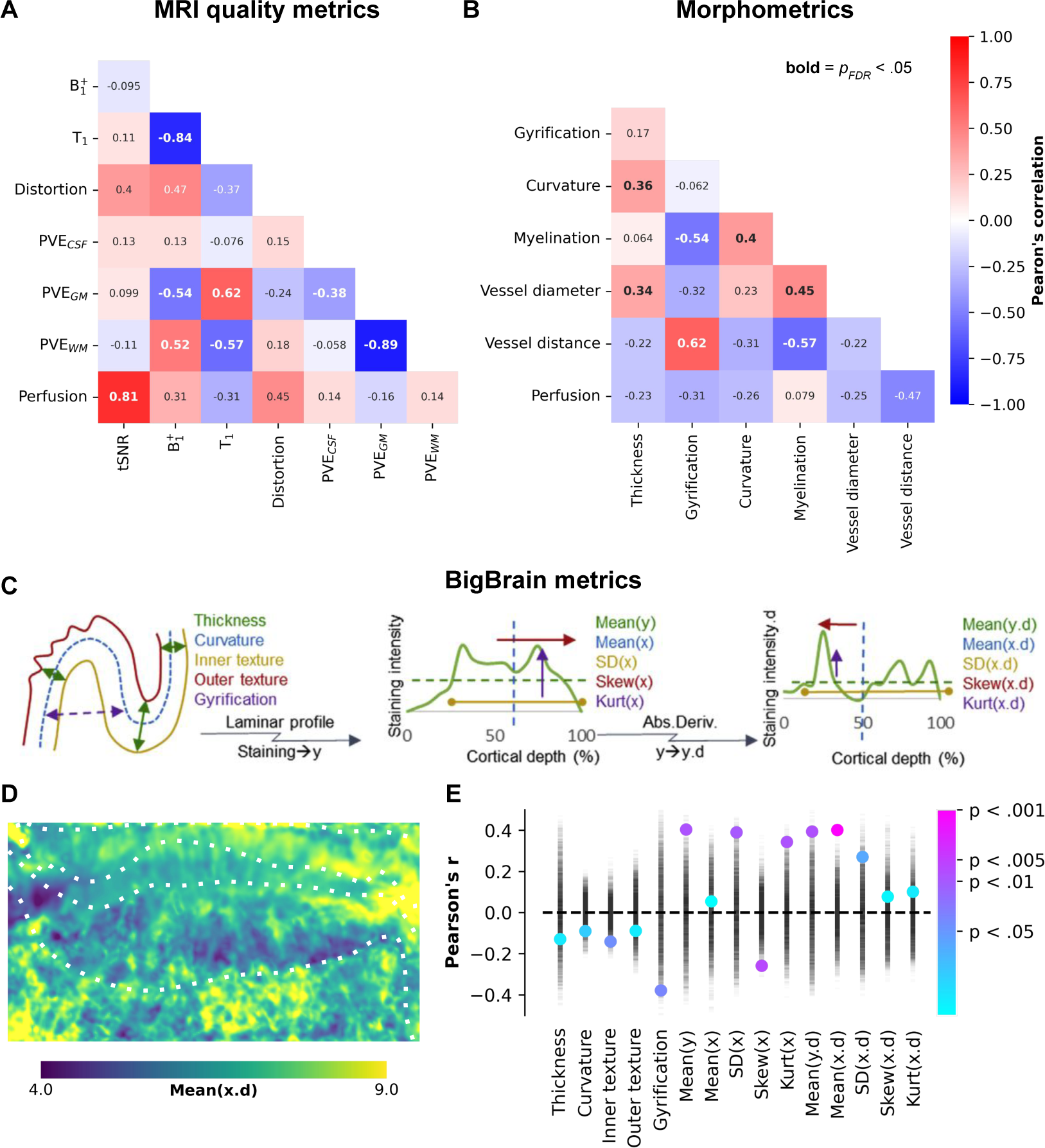
Between-feature correlations. (A-B) Heatmaps depicting the correlations between different features (see Supplementary Figs. 2 and 3) on their vertex-wise averages, with corresponding Pearson’s correlation coefficients annotated. Significant correlations, after correcting for spatial autocorrelation and multiple comparisons, are indicated by bold annotations. Panels (C-E) illustrate the correlations between perfusion and various hippocampal morphometric and cell density measures derived from BigBrain. In panel (E), the point plot displays permuted Pearson’s correlation coefficients represented by semi-transparent black markers, which were used to calculate color-coded significance levels.

Regarding the two macrovascular features, we found the strongest correlation between perfusion measures and distance to vessels (Pearson’s *r* = −.47, *p_roll_* = .06), indicating that regions further away from vessels tend to exhibit lower perfusion (see Figs. 1A and 3E). The impact of vessel diameter on measured perfusion was relatively smaller, consistent with the notion that smaller vessels, which are more relevant to tissue perfusion, are typically in closer proximity to the hippocampal grey matter (Fig. 3D).

One hypothesis for the relatively modest correlation between perfusion and the other (morphometric) features might be that perfusion levels are weighted stronger towards local differences in metabolic demand. To explore this further, we repeated the correlation analyses using cytoarchitectonic features derived from the BigBrain model (Fig. 4C)^31,34^. These cytoarchitectonic measures provide insights into the distribution of cell bodies within the hippocampal grey matter across its three axes (anterior-posterior, proximal-distal, and cortical depths) and can serve as proxies for variability in metabolic activity (i.e., heightened activity and functional requirements). Fig. 4D shows an example of one of these features, linked to the center of cell body density mass along the cortical depth direction (i.e, derivative of mean *X*). Among all tested features, perfusion appears most strongly (i.e., significantly, *p_roll_ <* .05) correlated with the hippocampus’ cytoarchitectonic rather than its morphometry aspects (Fig. 4E), suggesting a stronger dependence of perfusion on laminar features and possible associations with metabolic demand.

## Discussion

Emerging research suggests that certain subfields exhibit selective vulnerability to different types of disorders or conditions^14^. Previous studies have shown that part of this specificity can be ascribed to differences in the molecular profiles across subfields^35–37^, such as expression of NMDA and mineralocorticoid receptors and the flexibility to deal with metabolic insults (e.g., hypoxia, ischaemia and reductions in the level of circulating hormones)^38,39^. However, these factors provide only partial substrates for the selective vulnerability and it is hypothesized that non-molecular factors play a role as well, such as differences in cytoarchitecture^3^ and physiology (e.g., spiking rate)^40^. It is therefore likely that regional differences in metabolic demand due to their unique cellular configurations and activity render hippocampal subfields differently perfused by blood. As such, characterization of their perfusion will provide important insight to further our understanding of the hippocampus’ functioning in health and in disease.

### Perfusion in the human hippocampus

Alterations in hippocampal perfusion have been observed in various diseases such as Alzheimer’s disease^41^, temporal lobe epilepsy^42^ and schizophrenia^43^. However, these findings have primarily relied on imaging techniques such as positron emission tomography (PET), single-photon emission computed tomography (SPECT), or ASL with limited spatial resolutions (i.e., *>*2.5mm isotropic) and, hence, did not allow quantification of perfusion differences at a subfield level. The quantification of hippocampal subfield-specific perfusion requires optimized imaging acquisition and analysis strategies. Therefore, the objective of this study was to establish an imaging framework that enables users to accurately assess variations in hippocampal perfusion among its subfields. Here we show that it is possible to acquire robust perfusion-weighted data with consistent slab positioning across all subjects (Fig. S5A/B) for high resolution (1.5 mm isotropic) perfusion quantification using ASL at 7T. The perfusion measures, averaged across our cohort, fell within the expected physiological range in healthy humans (Fig. S5C)^44,45^. For reference, the perfusion in the visual cortex were V1: 58.24 *±* 15.68 ml/100 g/min, V2: 44.42 *±* 10.91 ml/100 g/min. The quantitative perfusion values observed in the hippocampus, although lower than those in V1 and V2, are unlikely to be artifactual based on the robustness of our data. Instead, they are likely attributed to the relatively lower microvascular density, which serves as the source of our perfusion signal, in the hippocampus compared to neocortical tissue (like V1 and V2)^46,47^. We demonstrate for the first time, that there are clear, measurable differences between subfields. Most strikingly, CA1 appears to be characterized by the lowest perfusion among hippocampal subfields, which is in line with previous *in vivo* and ex vivo indices of microvascular density in animals^48^ and humans^24,49^. Whilst characterized by a lower microvascular density and blood flow, CA1 is not necessarily characterized by a difference in activity due to the prominent role of its (mostly pyramidal) neurons in hippocampal structure and function^50^. This thus renders CA1 particularly vulnerable in case of metabolic insults and confirms its observed higher susceptibility across several diseases^14,51^. Furthermore, our stability analyses have demonstrated that the observed perfusion pattern stabilizes quickly and can be reliably detected with a relatively small sample size of six subjects and a total ASL scan time of only five minutes per subject (*∼*50 perfusion-weighted images). These findings indicate that a general-purpose high-resolution ASL protocol at 7T, as employed in this study, is capable of providing sufficient perfusion information in medial-inferior cortical regions like the hippocampus. Therefore, it suggests that specific optimization tailored to each region is not necessarily required^44,52^. However, it is worth noting that the above recommendation was based on data obtained from healthy and experienced control subjects. For researchers, particularly those investigating hippocampal perfusion in clinical populations, we advise acquiring as much ASL data as feasible within the available scan time to ensure comprehensive analysis and accurate interpretation of the findings.

### Vascularisation and its impact on hippocampal perfusion

Based on the aforementioned considerations, one may reasonably attribute the relatively diminished perfusion observed in the CA1 region and its heightened susceptibility to disease to its comparatively lower microvascular density. However, it is likely that the observed differences in perfusion across the hippocampal subfields were not only impacted by the density of small blood vessels but also by their proximity to the nearby macrovasculature. This intricate network of arteries and vessels supplies oxygen and nutrients to the hippocampal tissue and supports its metabolic demand and proper functioning^53^. The two primary arteries involved in hippocampal perfusion are the posterior cerebral arteries (PCAs) and the anterior choroidal arteries (ACHAs). However, the vasculature of the brain is highly interconnected, and there may be additional contributions from other arteries to hippocampal perfusion, including the hippocampal branches of the middle cerebral arteries (MCAs)^53^. We employed TOF-MRA to map subject-specific vessel branching patterns around the hippocampus *in vivo*, generating reconstructions consistent with previous descriptions of hippocampal vascularisation^22,23,53^. Our reconstructions, along with joint analyses of perfusion estimates, suggests that the subiculum’s perfusion is most likely provided by collateral branches of the PCA’s P2 segment — a vessel that runs parallel to the anterior-posterior hippocampal axis and exhibits a larger diameter. Most importantly, these results suggest that the lower perfusion in CA1 might indeed be partly ascribed to its further distance from the macrovasculature, especially towards the hippocampal head. While it is possible that increased partial voluming of perfusion-weighted signals between arteries and the subiculum, CA2, CA3, and CA4/DG might have artificially elevated their perfusion estimates, the dominance of gray matter tissue contributions observed in the perfusion analyses decreases the likelihood of this scenario^54^. Combining measurements of both distance and diameter demonstrates that the relationship between mean perfusion and distance is strongest when considering smaller vessels (i.e., *<*2mm), whereas increased variability in perfusion across subjects is more closely associated with the closer proximity of relatively larger vessels (i.e., *>*2mm). These integrative analyses collectively indicate that macrovascular structures likely influence the measured perfusion pattern and introduce variability in hippocampal perfusion measurements among subjects. Therefore, it is advisable for future work to compare hippocampal perfusion maps between groups of subjects characterized by different hippocampal vascularisation patterns to gain insights into the observed differences, particularly in the context of disease^22,55^.

### Methodological aspects of quantifying hippocampal perfusion

While we have successfully demonstrated the feasibility of reliably characterizing hippocampal perfusion, it remains a challenging task that necessitates certain expertise to ensure high-quality data. In this study, we implemented a multi-modal, multi-resolution acquisition protocol for 7T MRI. The use of 7T MRI offers improved image quality compared to 3T MRI, thanks to increased SNR^28^ and potential enhancements in spatial resolution. This enhancement allows for better anatomical delineation of hippocampal subregions^56^ and improved sensitivity to perfusion differences^57^. However, the inclusion of scans with small field-of-view and different orientations introduced an additional challenge in terms of data integration. This challenge becomes evident when overlaying the slab positioning for the various acquisitions (refer to Fig. S6). Nevertheless, not all of these scans are equally critical. In the following discussion, we address this aspect and propose a set of minimal requirements to be considered when conducting hippocampal perfusion imaging, ensuring feasibility and data quality.

For anatomical imaging, we recommend acquiring at least an MP2RAGE image^58^ and a B^+^ map (e.g., using the Sa2RAGE sequence^59^) to improve hippocampal T_1_ quantification^60,61^, which subsequently enhances the precision of voxel-wise perfusion estimates^62^. Consistency in subfield labels and harmonization across subjects are crucial to maximize the spatial specificity of hippocampal perfusion maps^4^. Therefore, in this study, we opted to acquire additional T_2_-weighted images to extract hippocampal surfaces and perform subfield parcellation using the HippUnfold analysis suite^29^. The Hippocampal Subfields group suggests the use of T_2_-weighted images for manual segmentation of the hippocampus because of their optimal contrast between hippocampal gray matter and stratum radiatum and lacunosum-moleculare (SLRM) tissue^17,63^. Fig. S5 D-G provides an example of manual segmentation and corresponding surface representation for the left and right hemispheres of a single subject. While T_2_-weighted images are generally preferred, recent advancements in HippUnfold enable precise and automatic segmentation even when only T_1_-weighted data are available^29^. Furthermore, although the CA4 field and DG are distinct anatomical entities^3^, they were combined into a single label due to their limited size. However, in the most recent releases of HippUnfold (v1.0.0 and newer), the DG is modelled as a separate surface to increase specificity.

Furthermore, we would like to emphasise several aspects regarding our perfusion imaging protocol. We optimised the high-resolution ASL protocol to be acquired in approximately five minutes per run. This optimisation ensured robustness against subject motion during scanning and minimised data loss, which is particularly crucial for clinical applications. Additionally, ASL-based perfusion imaging is a B^+^ sensitive technique and therefore challenging to acquire at 7T due to its transmit field inhomogeneities, especially towards the lower part of the brain (e.g., inferior frontal and temporal lobes)^64^. This consideration is important to note when transitioning from 3T to 7T for perfusion imaging. To address this, we employed dielectric pads^65,66^ and an optimised inversion (TR-FOCI^67^) pulse to achieve higher labelling efficiency (i.e., *α*=0.95) in the hippocampal region (Fig. S2B) and the adjacent vasculature (Fig. S4B) for all subjects^27,66^. Lastly and more generally, reducing geometric distortions, high-spatial resolution and isotropic voxels are crucial to reduce partial voluming effects and thereby, improve the perfusion CNR^68^. Some ASL protocols employ 3D-GRASE readouts to obtain relatively higher SNR^69^ but they come at the cost of increased blurring in the z-direction^70^ as well as higher SAR at ultra-high field strengths. Alternatively, ASL with spiral^71^ and 3D-EPI^72^ readouts have shown promise to enable high-resolution, SAR efficient perfusion imaging at ultra-high fields. In this study, we did not sought out to optimise the PLD parameter for hippocampal imaging in particular due to our cohort consisting of young, healthy participants and to prevent erroneous estimation of hippocampal grey matter perfusion^73^. However, this should be considered when imaging other cohorts such healthy elderly or patients as the longer arrival times necessitate increasing PLD to obtain robust perfusion^74^.

### Concluding remarks

By quantifying blood flow across hippocampal subfields, we can gain a better understanding of the normal patterns of perfusion and how they relate to the specific functions associated with each subfield. Here we presented and validated a 7T MRI imaging framework that allows *in vivo* characterization of perfusion differences across the hippocampus. Our hippocampal perfusion map can serve as a baseline for comparison with diseased states where it might possibly allow for early detection and/or assessment of disease progression in individuals with hippocampal-related disorders. Diseases that cause even modest reductions in hippocampal blood flow, potentially due to capillary rarefaction, hyperconstriction and inward remodeling of hippocampal arterioles, would likely have a tremendous impact on neuronal function, memory and cognition^75^.

## Methods

Eleven healthy volunteers (mean age 26±3.2 years, 5 males) participated in this study after having provided written informed consent. The study was approved by the Ethics Review Committee Psychology and Neuroscience (ERCPN) at the Faculty of Psychology and Neuroscience, Maastricht University, The Netherlands, and all procedures followed the principles expressed in the Declaration of Helsinki.

### Data acquisition

All data were acquired on a Siemens Magnetom 7T scanner (Siemens Healthineers, Erlangen, Germany) with an SC72 whole-body gradient system capable of maximum gradient amplitude of 70 mT/m, maximum slew rate of 200 T/m/s using a 1Tx/32Rx phased array head coil (Nova Medical, USA) housed at Scannexus B.V., Maastricht, The Netherlands. The participant preparatory and positioning procedure followed the protocol previously described in^27,57,72^. Briefly put, the centre of the eyes were used as the iso-centre reference (instead of the eyebrows, as is typically done), supplemental cushions were provided to the participants under the neck, to ensure that the large feeding arteries to the brain were as close to parallel to the B_0_ as possible. In addition, two 13×13×0.5 cm^3^ high-permittivity dielectric pads containing a 2.8:1 solution of calcium titanate (CaTiO_3_) in heavy water (D_2_O) by weight^76^ were placed on either side of the neck to improve the B^+^ (therefore, labelling) efficiency at 7T^65^. In 6 participants, a third dielectric pad was placed over the participant’s right lateral side to reduce the impact of the hemispheric asymmetry of the coil’s inherent B^+^ profile^66^.

#### Anatomical data

A whole-brain 3D-MP2RAGE^58^ dataset at 1.0 mm isotropic resolution was acquired first and used to inform slice positioning during the rest of the session. A 3D-Sa2RAGE^59^ dataset at 2 mm isotropic was acquired to facilitate B^+^ correction of the T_1_ maps^77^. At least three repetitions of an ultra-high-resolution 0.4 mm in-plane resolution T_2_-weighted 2D-TSE^78^ were acquired using oblique coronal slices positioned to cover the entire hippocampal complex bilaterally. Two ultra-high resolution 0.5 mm isotropic 3D-MP2RAGE scans with a partial coverage (entire hippocampal region axially) were acquired. Due to SAR constraints, 2D-TSE scans could not be acquired consecutively, the 0.5 mm 3D-MP2RAGE scans were interspersed between the 2D-TSE scans for time efficiency. Finally, two repetitions of ultra-high resolution 0.5 mm isotropic 3D multi-slab time-of-flight^79–81^ MR angiograms were acquired (3D-TOF-MRA). Complete sequence details are tabulated in Table 1.

**Table 1.** MRI acquisition details. See Supplementary Fig. 6 for a schematic of the scanning order and positioning of the imaging slabs.

|  | Anatomy |  |  | Perfusion | Vasculature |
| --- | --- | --- | --- | --- | --- |
| Parameter | MP2RAGE |  | TSE T<sub>2</sub>w | ASL | TOF-MRA |
| *TR (ms)* | 6000 | 6000 | 9000 | 2861 | 15 |
| *TE (ms)* | 1.88 | 3.98 | 105 | 14 | 3.59 |
| *TI<sub>1</sub>/TI<sub>2</sub> (ms)* | 800/2750 | 983/2940 |  | 700/1800 |  |
| *FA<sub>1</sub>/FA<sub>2</sub> (°)* | 4/5 | 6/7 | 132 | 70 | 15 |
| *GRAPPA* | 4 (A-P) | 2 (A-P) | 2 (F-H) | 3 | 3 (F-H) |
| *No. of slices* | 192 | 72 | 50 | 32 | 220 |
| *Slice direction* | Sagittal | Sagittal | Coronal | Sagittal | Coronal |
| *Field of view (mm)* | 256×256 | 184×184 | 192×192 | 192×192 | 210×210 |
| *Matrix size* | 256×256×192 | 368×368×144 | 512×512×50 | 128×128× | 448×448×440 |
| *Resolution (mm)* | 1×1×1 | 0.5×0.5×0.5 | 0.4×0.4×1 | 1.5×1.5×1.5 | 0.5×0.5×0.5 |
| *Phase partial Fourier* | 6/8 | Off |  | 6/8 | 6/8 |
| *Slice partial Fourier* | Off | 6/8 |  |  | 6/8 |
| *Bandwidth (Hz/px)* | 250 | 140 | 90 | 1698 | 203 |
| *Acquisition time (m:s)* | 7:14 | 5:26 | 4:41 | 4:49 | 5:50 |
| *Number of runs* | 1 | 2 | 3 | 8 | 2 |
| *Total acquisition time* | 1:20:21 (h:m:s) |  |  |  |  |

#### Perfusion data

Perfusion data was acquired at 1.5 mm isotropic resolution using a Pulsed Arterial Spin Labelling (PASL) sequence^82^ employing a FAIR^83^ QUIPSS II^84^ labelling scheme with a 2D-EPI readout. For each participant, eight consecutive runs of 50 control-label repeats (i.e., 100 volumes) were acquired with each run lasting *±*5 min. An equilibrium magnetisation (M_0_) image was acquired using the same PASL sequence and 2D-EPI readout, but with no magnetisation preparation and the TR increased to 20 s. A second M_0_ image was acquired immediately after with the opposite phase-encoding direction for distortion correction.

### Anatomical data processing

All stages of data processing and registrations were subject to careful visual inspection for quality control.

#### TSE

The TSE runs were first resampled to 0.3 mm isotropic resolution using a 5*^th^* order B-Spline interpolation with ANTs’s *ResampleImage*^3^. A minimally deformed average TSE template was created from the 0.3 mm TSE datasets using ANTs’s *antsMultivariateTemplateConstruction2.sh* script^85^. This resampled 0.3 mm isotropic TSE template image was used for manual hippocampal segmentation and was defined as the final reference space for co-registering all other image modalities in the present study.

#### MP2RAGE

Signal from dielectric pads were first masked out^4^ of both whole-brain (1 mm^3^) and high-resolution (0.5 mm^3^) MP2RAGE datasets (forthwith referred in text using prefixes ‘wb-’ and ‘hires-’, respectively) following which they were corrected for transmit efficiency (B^+^) inhomogeneities using a separately acquired Sa2RAGE B^+^ map^59^ in line with^77^, and following the code and procedure provided by^5,60^. The B^+^ corrected MP2RAGE UNI images were then pre-processed using *presurfer*^686^. The cleaned wb-UNI image was used as input using the default recon-all pipeline for cortical segmentation and surface reconstruction in Freesurfer 7.1.1^87, 7^. The cleaned hires-UNI, and the B^+^ corrected hires-UNI images and hires-T_1_ maps were resampled to 0.3 mm isotropic resolution using a 5*^th^* order B-Spline interpolation with ANTs’s *ResampleImage*. A minimally deformed average template image was created using ANTs’s *antsMultivariateTemplateConstruction2.sh* script and the average B^+^ corrected hires-UNI image and hires-T_1_ map were used in further analyses.

#### TOF-MRA

The TOF-MRA MRA data were first resampled to 0.3 mm isotropic resolution using a 5*^th^* order B-Spline interpolation with ANTs’s *ResampleImage*. Next, the second run was co-registered to the first run by a rigid-body transformation using *greedy* with the Neighbourhood Cross Correlation (NCC) metric. Then, the estimated transformation matrix was converted to an ITK matrix using *c3d_affine_tool* and the second run was resampled using ANTs’s *antsApplyTransforms* and its Lanczos Windowed Sinc interpolator. An average TOF-MRA image was calculated using ANTs’s *AverageImages* and this average 0.3 mm isotropic TOF-MRA image was used for vascular segmentation.

### Perfusion data processing

First, the ‘blip-up’ and ‘blip-down’ M_0_ EPI datasets were rigidly realigned to their respective first volume in the timeseries using FSL’s *flirt* with the NMI cost function (normmi) and resampled using the spline interpolator. Then, a temporal mean was calculated from the realigned M_0_ timeseries. Next, a rigid-body registration was estimated from the blip-down (moving image) to the blip-up (fixed image) using FSL’s *flirt*. The blip-up and registered blip-down M_0_ image were combined into a 4D file using FSL’s *fslmerge* and the phase-encoding distortion correction was estimated using FSL’s *topup*^33^.

All ASL images were rigidly motion-corrected using the blip-up M_0_ as a reference space using an iterative implementation of FSL’s *flirt*. Motion matrices and phase-encoding distortion estimate were combined into a warp using FSL’s *convertwarp*. All ASL runs were corrected for motion and phase-encoding distortions using a single resampling step using FSL’s *applywarp* and spline interpolation. Perfusion-weighted images (PWI) were calculated from the ASL timeseries datasets using the surround-subtraction approach^88,89^ as implemented in FSL’s *asl_file*. Perfusion temporal signal-to-noise (tSNR) map was calculated by dividing the PWI temporal mean by the PWI temporal standard deviation. Perfusion quantification was carried out in native space using *oxasl*^8^ using the PASL model^44^. The following parameters were modified as per our acquisition scheme (inversion efficiency = 0.95,^57^) and the field-strength (T_1_,blood = 2.2 s,^90^; subject-wise T_1_ image was provided using *–t1img*^32,77,91^).

### Registration to 0.3 mm TSE space

#### Anatomical data

The wb-UNI (moving image) was co-registered to the hires-UNI (fixed image) by a rigid-body transformation using *greedy*,^992^ with the Normalised Mutual Information (NMI) cost function. The registrations was visually inspected for quality control. The estimated transformation was applied using *greedy* and its LABEL interpolator to resample the WM segmentation from Freesurfer to the hires-MP2RAGE space. The estimated transformation matrix was converted to an FSL compatible matrix (‘wb2hires’) using *c3d_affine_tool*^10^.

The transformation between the hires-MP2RAGE and TSE datasets was estimated in two stages. First, a rigid-body registration was estimated from the TSE (moving image) to the hires-UNI (fixed image) using *greedy* with a Normalised Mutual Information (NMI) metric. Then, *c3d_affine_tool* was used to convert this c3d matrix to an FSL matrix. The second stage involved use of the boundary-based registration (BBR^93^) cost function as implemented in FSL’s *flirt* together with the initialisation matrix from the first stage to register the TSE (moving image) to the hires-UNI (fixed image) in a robust manner. The registrations was visually inspected for quality control. The resulting transformation matrix (i.e. ‘tse2hires’) was inverted using FSL’s *convert_xfm* to obtain the hires-MP2RAGE to TSE transformation (‘hires2tse’). Finally, this transformation matrix was applied using the spline interpolator in FSL’s *flirt* to the hires-T_1_ map to transform it to the TSE space.

#### TOF-MRA data

The TOF-MRA (moving image) was co-registered to the TSE (fixed image) by a rigid-body transformation using *greedy* with the NCC metric. The registration was visually inspected for quality control. Then, the estimated transformation matrix was converted to an ITK compatible matrix using *c3d_affine_tool*. Finally, the TOF-MRA was resampled using ANTs’s *antsApplyTransforms* and its Lanczos Windowed Sinc interpolator.

#### Perfusion data

The registration strategy to transform the ASL data into the TSE was as follows. First, a rigid-body transformation matrix was estimated from the ASL data (moving image) to the wb-UNI image using FSL’s *flirt* with the BBR cost function (‘asl2wb’). The ‘asl2wb’ and ‘wb2hires’ (estimated previously) transformation matrices were concatenated using FSL’s *convert_xfm* to obtain the ‘asl2hires’ transformation matrix, which is the affine transformation from the ASL native space to the hires-UNI space. Second, the ‘asl2hires’ and ‘hires2tse’ (estimated previously) transformation matrices were concatenated to obtain the ‘asl2tse’ transformation matrix, which is the affine transformation from the ASL space to the 0.3 mm TSE reference space. All derivatives from the ASL data such as perfusion and perfusion tSNR were transformed from their native space to the TSE space using a single resampling step using FSL’s *flirt* and its spline interpolator by applying the final ‘asl2tse’ transformation matrix. The registration quality was visually inspected at every stage of the transformation including all the intermediate steps.

### Hippocampus and subfield segmentation

The 0.3 mm isotropic average TSE data were used to manually segment the hippocampus for each subject (Supplemetnary Fig. 5D). In the average TSE data, the contrast between the stratum radiatum and lacunosum-moleculare (SLRM), or ‘dark band’, and the neighbouring hippocampal GM tissue is improved and was essential to facilitate manual segmentation. First, individual masks for both SLRM and GM tissues were created semi-automatically using the active contour segmentation mode in ITK-SNAP v3.8.0^94^ and were manually edited following the recommendations in^34^. Additionally, several ‘boundary’ labels were added to encode for the anterior-posterior (A-P), proximal-distal (P-D) and inner-outer (I-O) axes (Supplemetnary Fig. 5E).

Following the manual segmentation, each hippocampus was unfolded using the snakemake^95^ implementation of our in-house developed hippocampal unfolding tool (Fig. S5F-G)^29^. In brief, this method entails the following steps: (i) alignment of the subject-specific T_2_w image and its manual segmentation to the coronal oblique atlas space, (ii) imposing coordinates along the A-P, P-D and I-O dimensions onto the hippocampal GM by solving the Laplace equation, (iii) extracting inner, mid-thickness and outer GM surfaces whilst ensuring one-to-one vertex correspondence between them, and (iv) estimating the native-to-unfolded space transformation to analyse data in a common 2D plane. A detailed description of the unfolding algorithm can be found in the original work^34^ and online documentation^11^. All the surface-based output was generated within the GIfTI framework to allow easy manipulation, volume-to-surface mapping (see following sections) and visualization using Connectome Workbench^96^. Exploration of the manual segmentations and HippUnfold output is possible using the HTML visualization notebooks provided in the online code repository.

### Vascular segmentation and reconstruction

The average 0.3 mm TOF-MRA data were used to identify macrovascular structures within the vicinity of the hippocampus. First, the TOF-MRA image was spatially filtered using non-linear anisotropic diffusion^97,98^ by exploiting the structure tensor field derived from the images as implemented in Segmentator v1.6.0^99, 12^. This preserves the boundaries between vessels and brain tissue while reducing intra-tissue class image noise. Next steps were carried out in MeVisLab v3.3^100, 13^. First, vessel-like structures were extracted from the ‘smoothed’ TOF-MRA image for 3D reconstruction. Then, for each subject and hemisphere, the input image was rescaled to range between 0-100 a. u. (i.e., to match intensity ranges across subjects), then thresholded to increase the contrast between vessels and background (i.e., GM and WM, image intensity *<* 10, a. u.) tissue and finally used to manually define *±*150 seeding points to segment connected vessels. This ensures that all voxels connected in the x, y or z direction with a seed point, and within the specified intensity range will be segmented. Here, the lower threshold was optimised for each subject based on manual inspection of the vascular tree after its automatic 3D reconstruction. This was achieved by (a) extracting the vessels’ skeleton based on the centerline of the binary segmentation label, (b) transforming the skeleton into a graph to encode geometrical and structural shape properties so as to allow (c) the decoding of the graph properties into an polygonal surface of the vascular tree for 3D visualization^101^. Finally, surfaces were transformed to voxel-wise representations and skeleton graphs saved as an XML file for network reconstruction and analyses.

### Data integration and visualization

Subsequently, the hippocampal mid-thickness surface, ASL and TOF-MRA output maps were combined to assess their relationship. First, for each mid-thickness vertex, distance (in mm) to the most nearby vessel structure was calculated by taking the minimum euclidean distance to all vessel centreline voxels minus their radius^102^. These, as well as the respective vessel diameters, were then exported as GIfTI metric files using nibabel v3.2.0^103^. Second, for the co-registered imaging data, Connectome Workbench’s *-volume-to-surface-mapping* command-line tool was used to sample along the hippocampal GM midthickness vertices, hereby constraining the mapping algorithm to only include voxels that were labeled as GM and found between the inner and outer GM surfaces. As such, each vertex’s value represents a weighted average of the voxels along the IO dimension with lower weights for voxels positioned more distal to the mid-thickness surface.

Additionally, we developed a Python-based framework for network-based analyses of the vasculature’s structure. Skeleton graph XML files are parsed to define segment type (start, termination, branchpoint or skeleton) by examining the degree of connectivity, as well as connecting edges using the NetworkX package^104^. Each edge represents a physical connection between two nodes of type (i) start–skeleton, (ii) bifurcation–skeleton, (iii) skeleton–skeleton or (iv) skeleton–end with properties defining length, diameter, volume and surface area. Nodes and edges are used to construct a network for extraction of the shortest path from a given node to the rest of the vascular tree, as well as to compute different network characteristics (e.g., connected components, lowest common ancestors). Finally, vascular networks can be visualized and inspected interactively using implementation of the plotly interface. Individual MeVisLab workflows, output files and visualization notebooks for each subject and hemisphere can be found in the online repository.

### Statistical analyses

Statistical analyses were performed using the *pingouin* Python package^105^. The non-parametric Friedman’s test for repeated measures analyses of variance was used to assess differences across subfields. In case of a significant subfield effect, the Wilcoxon signed rank-test was applied for pairwise-comparisons, correcting for multiple comparison using the Benjamini-Hochberg false-discovery rate (FDR*_BH_*) method. The Pearson’s correlation coefficient was used to assess correlations among hippocampal surface maps (e.g., perfusion vs. T_1_w/T_2_w), while controlling for spatial autocorrelation^106^ using ‘roll’-based permutation testing (*p_roll_*) as well as multiple comparisons using FDR*_BH_* correction when constructing the correlation heatmaps. Briefly, to generate null distributions, N=5000 permuted maps are generated by randomly shifting the 2D hippocampal maps across one or both axes using SciPy’s *shift* function and through rotation using their *rotate* function^107^. Here, extension of maps was ensured by wrapping around to the opposite edge. Significance was then determined based on the position of the empirical correlation coefficient with respect to the generated null distribution^6^.

## Footnotes

1 https://tinyurl.com/3z8czuy9/

2 https://github.com/royhaast/hippocampal_perfusion

3 https://github.com/ANTsX/ANTs

4 https://github.com/srikash/ants_deface_depad/blob/master/PadsOff

5 https://github.com/JosePMarques/MP2RAGE-related-scripts

6 https://github.com/srikash/presurfer

7 https://surfer.nmr.mgh.harvard.edu

8 https://github.com/physimals/oxasl

9 https://github.com/pyushkevich/greedy

10 https://github.com/pyushkevich/c3d

11 https://hippunfold.readthedocs.io

12 https://github.com/ofgulban/segmentator

13 https://www.mevislab.de

## Acknowledgements

We would like to thank the participants who agreed to take part in this study. Author A.R.K received support by the Canada Foundation for Innovation (CFI) John R. Evans Leaders Fund project (grant agreement no. 37427), a CIHR Project Grant (366062) and Canada Research Chairs (950-231964). Author B.P. has received partial funding from the Dutch Science Foundation NWO under grant VIDI-TTW (016-178-052), European Union’s Horizon 2020 research and innovation program (885876) and NIH (R01MH111444). Authors R.A.M.H and J.K. were supported by BrainsCAN and Natural Sciences and Engineering Research Council of Canada postdoctoral fellowships, respectively.

## Author contributions statement

R.H. and S.K. designed research; R.H., S.K., M.Y., performed research; R.H., S.K., J.D. and A.K. contributed new reagents/analytic tools; R.H. and S.K., analyzed data and wrote the paper. All authors reviewed the manuscript.

---

**Supplementary Figure 1.**
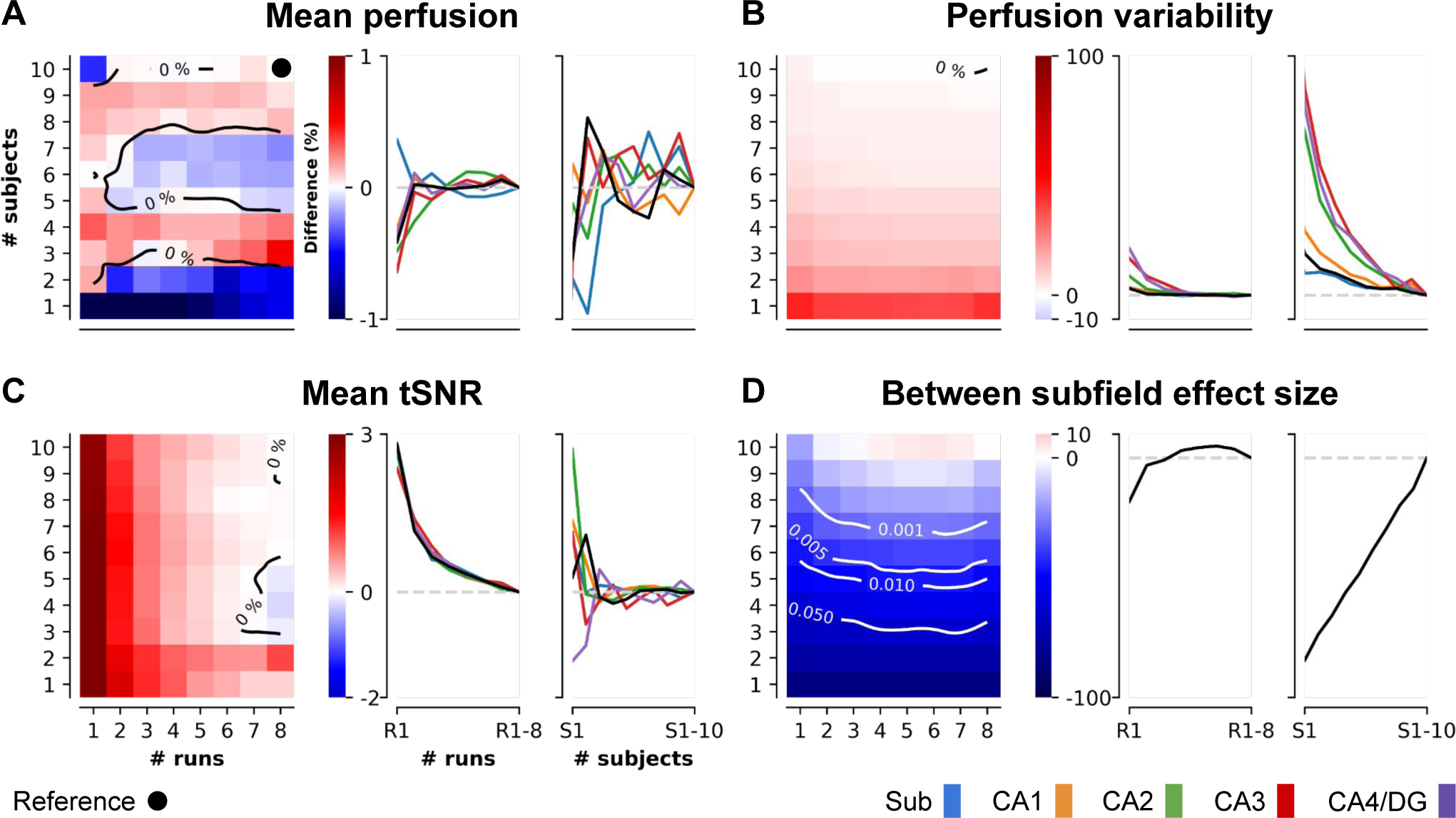
Bootstrap analysis. Evolution of (A) mean perfusion, (B) perfusion variability (coefficient of variation), (C) mean tSNR and (D) between subfields effect size using the median across N=100 bootstrap samples. For each metric, heatmaps depict the percentage difference with respect to the final estimates as function of number of included runs and subjects for global hippocampal estimates. Line plots show the impact of number of included runs (across all subjects) and subject (across all runs) on global and subfield-specific estimates. Superimposed contours indicate the 0% level for A, B and C and *p*-value thresholds for D.

**Supplementary Figure 2.**
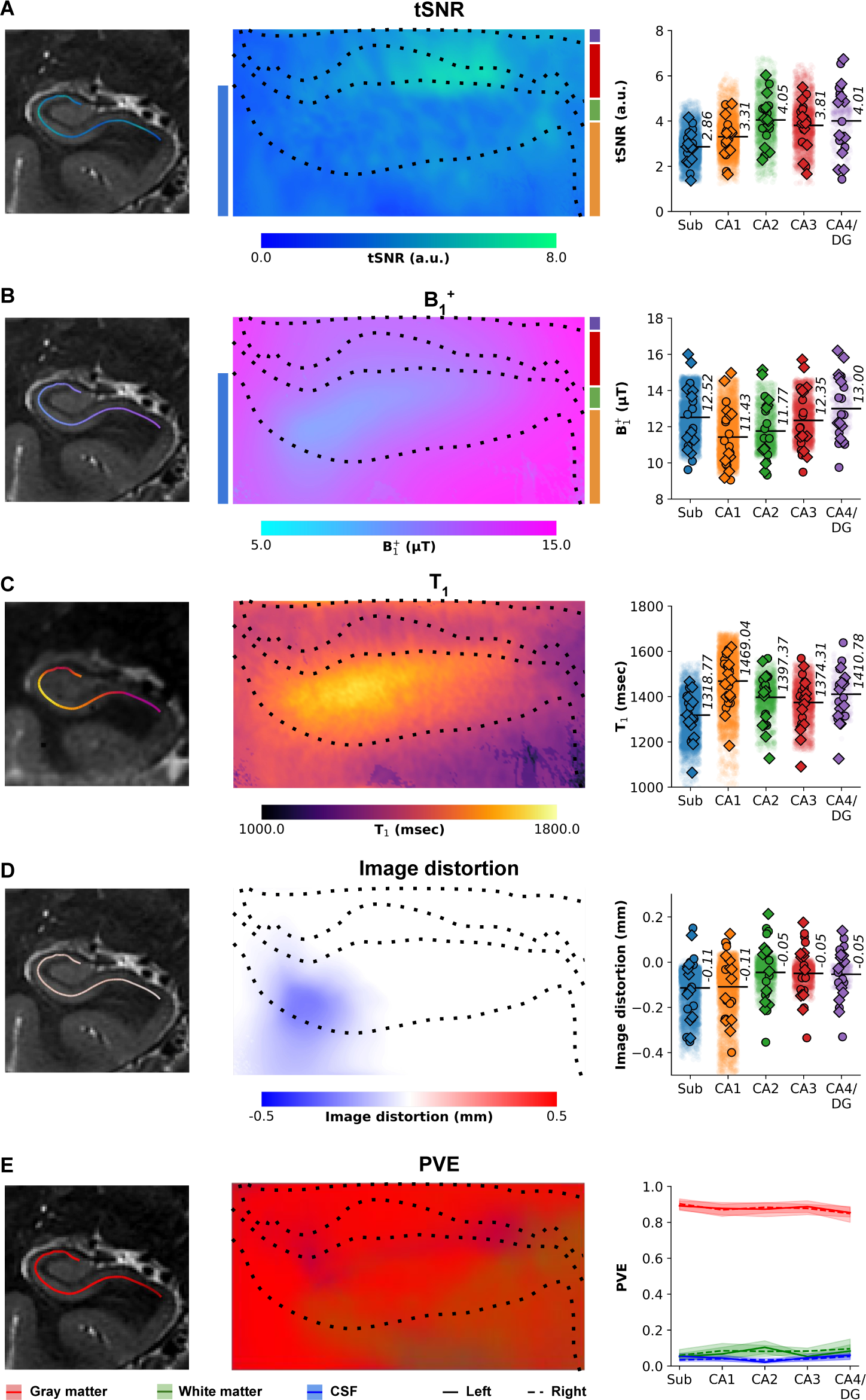
MRI quality metrics. Average (A) perfusion tSNR (a.u.), (B) B^+^ (*µ*T), (C) T_1_ (msec), (D) image distortion (mm) and (E) partial volume estimates (PVE) are mapped on the unfolded hippocampal surface. Dotted lines indicate subfield boundaries. The center plots show subfield averages for left (solid) and right (dashedline) hemispheres separately. Color-coded (as per subfield atlas overlaid on center images) subfield averages are shown for each subject and left (circles) and right (diamonds) hemisphere, as well as per vertex (i.e., averaged across subjects and hemispheres, semi-transparent dots, right plots). PVE estimates are displayed as line plots and color-coded based on tissue class.

**Supplementary Figure 3.**
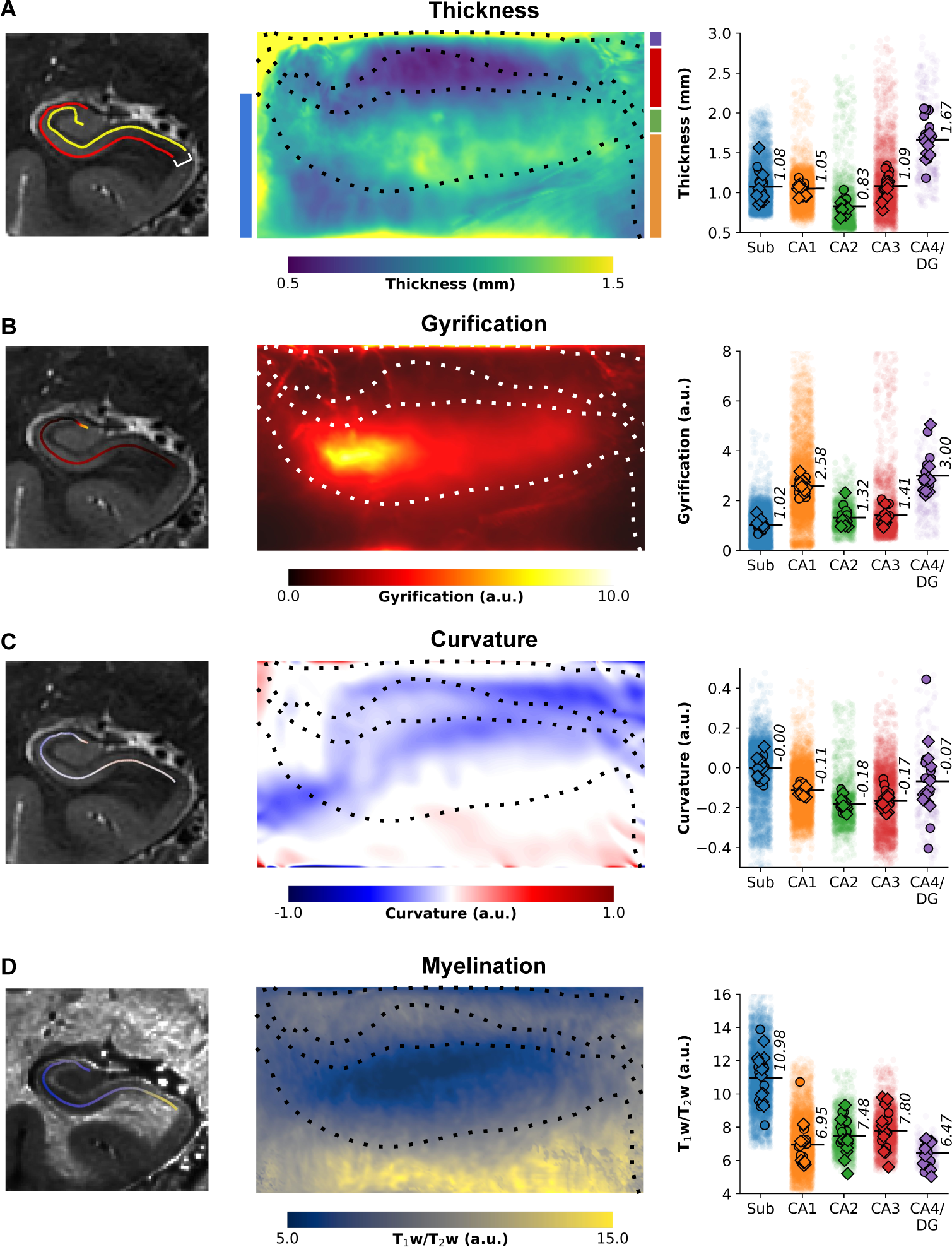
Morphometric hippocampal tissue properties. (A) Thickness (mm), (B) gyrification (a.u.), (C) curvature (a.u.) and (D) myelination (i.e., T_1_w/T_2_w, a.u.) are displayed for an example subject with color-coded surface outlines superimposed onto a coronal slice (left). Center images show the respective averages mapped on the unfolded hippocampal surface with dotted lines delineating subfield boundaries. Color-coded (as per subfield atlas overlaid on center images) subfield averages are shown for each subject and left (circles) and right (diamonds) hemisphere, as well as per vertex (i.e., averaged across subjects and hemispheres, semi-transparent dots, right plots).

**Supplementary Figure 4.**
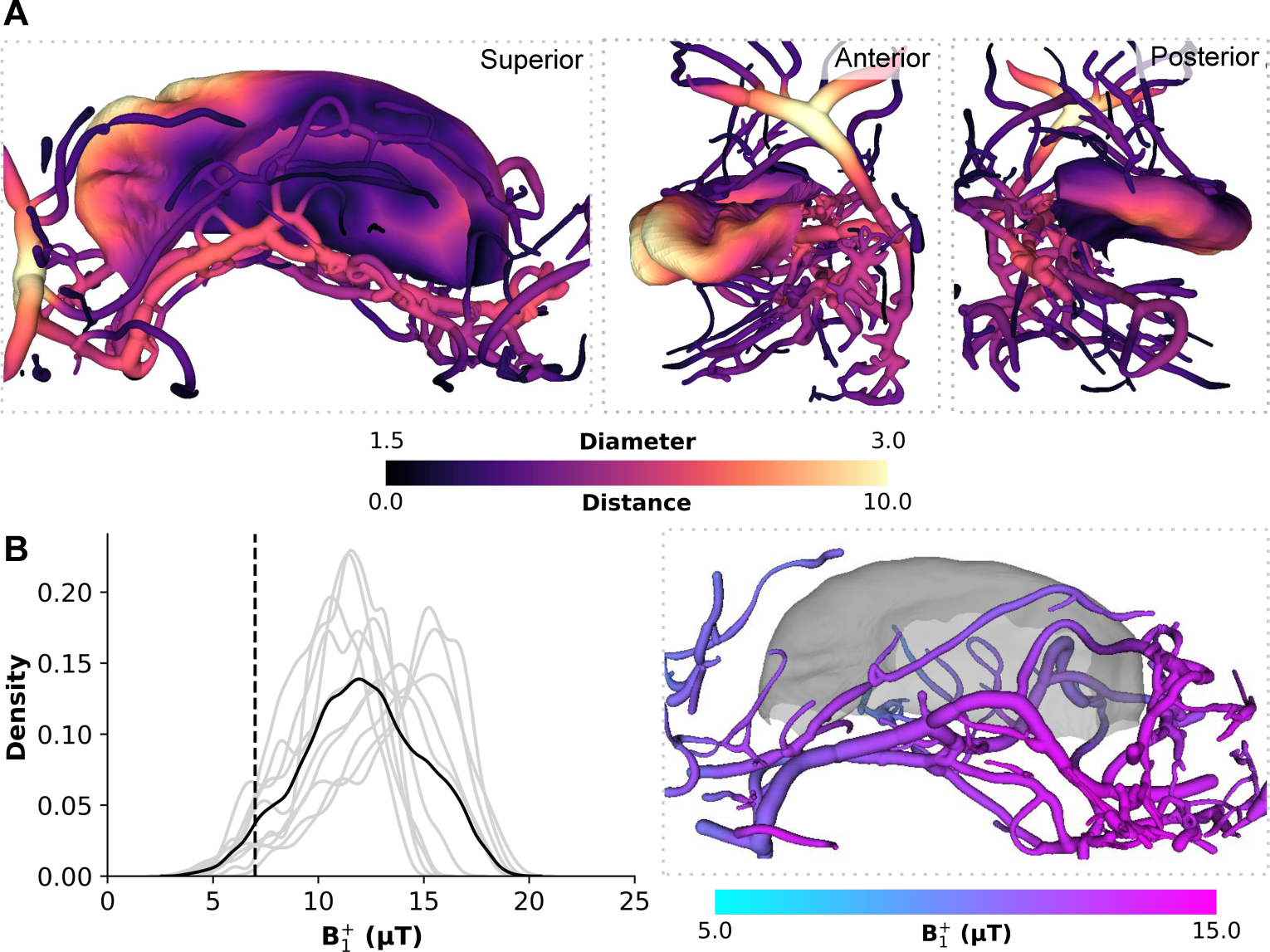
Hippocampal vasculature and grey matter projections. Example of a three-dimensional reconstruction of a subject’s macrovasculature near the right hippocampus color-coded for vessel diameter. Shortest distance between hippocampal vertices and the vessel tree is projected on the folded hippocampal surfaces. The colourmap on the vessels indicates local diameter (mm) while on the surface maps indicates the distance (mm) to closest vessel.

**Supplementary Figure 5.**
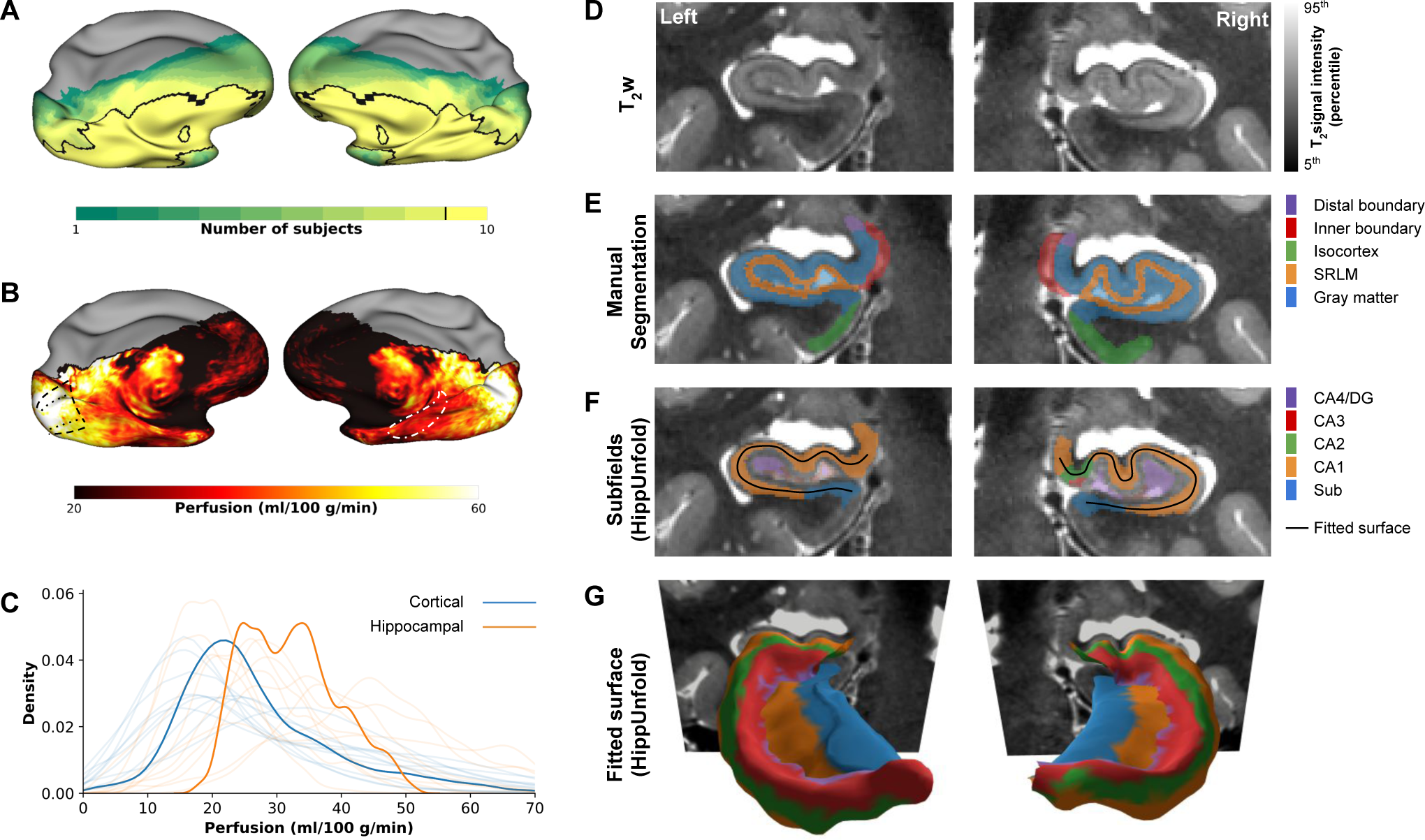
Hippocampal perfusion imaging and subfield segmentation. (A) Cortical projections of the vertex-wise coverage, and (B) average perfusion across subjects, (C) average perfusion distribution in cortex and hippocampus. (D) Example T_2_w data for a single subject’s left and right hippocampus, (E) manual segmentation of hippocampal tissue, (F-G) HippUnfold subfield labelling and fitted surface.

**Supplementary Figure 6.**
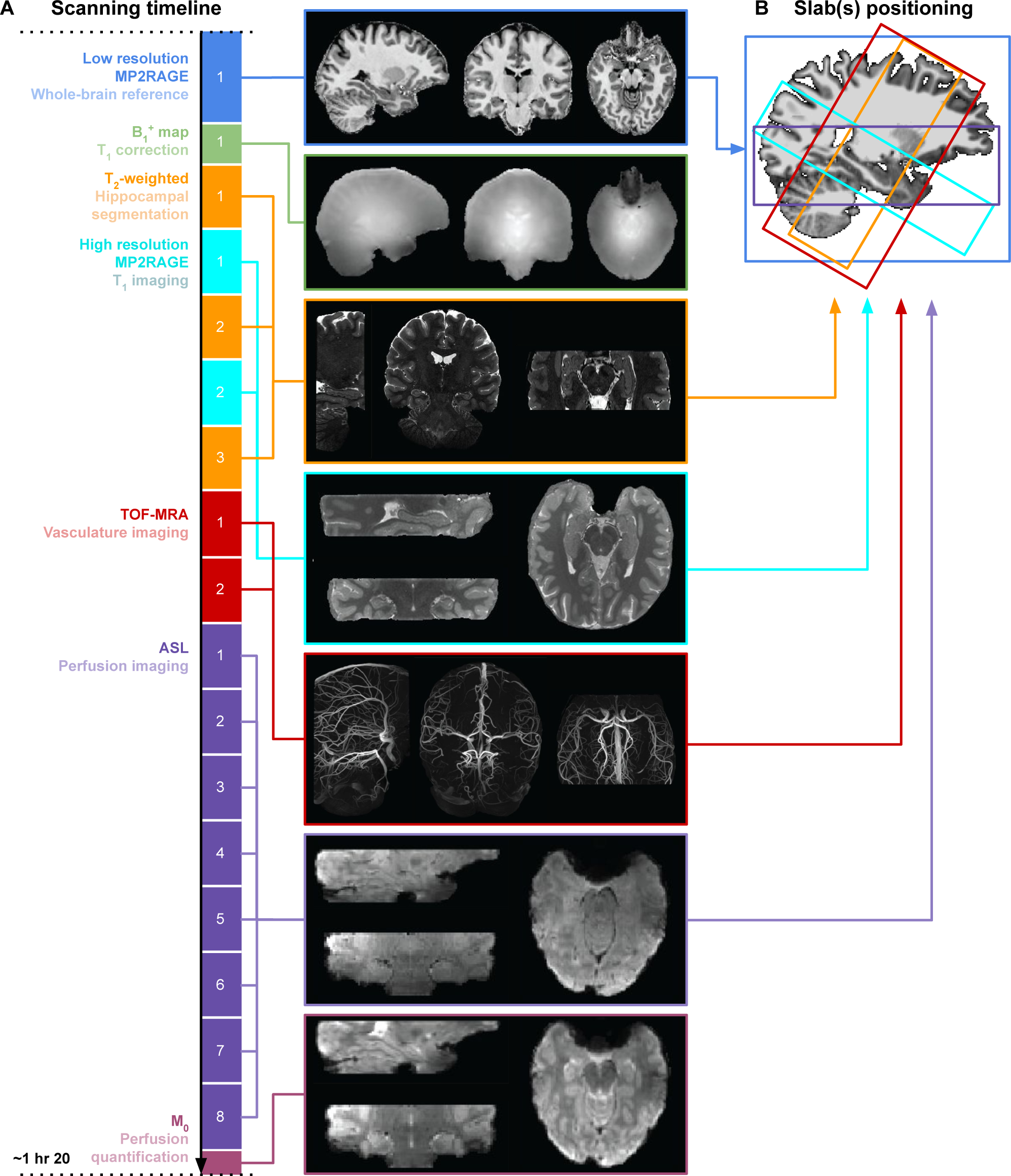
MRI modalities in the present study. (A) Scanning timeline showing the order of acquisitions colored by MRI modality. (B) Positioning of each MRI modality with respect to the whole-brain reference.

## References

1. Eichenbaum, H. Hippocampus: Cognitive Processes and Neural Representations that Underlie Declarative Memory. Neuron 44, 109–120, DOI: 10.1016/j.neuron.2004.08.028 (2004).

2. Squire, L. R., Stark, C. E. L. & Clark, R. E. The medial temporal lobe. Annu. Rev. Neurosci. 27, 279–306, DOI: 10.1146/annurev.neuro.27.070203.144130 (2004).

3. Ding, S.-L. & Van Hoesen, G. W. Organization and Detailed Parcellation of Human Hippocampal Head and Body Regions Based on a Combined Analysis of Cyto- and Chemoarchitecture. The J. Comp. Neurol. 523, 2233–2253, DOI: 10.1002/cne.23786 (2015).

4. DeKraker, J., Köhler, S. & Khan, A. R. Surface-based hippocampal subfield segmentation. Trends neurosciences 44, 856–863 (2021).

5. Genon, S., Bernhardt, B. C., La Joie, R., Amunts, K. & Eickhoff, S. B. The many dimensions of human hippocampal organization and (dys) function. Trends neurosciences 44, 977–989 (2021).

6. Karat, B. G., DeKraker, J., Hussain, U., Kohler, S. & Khan, A. R. Investigating the in vivo spatial distribution of hippocampal microstructure and macrostructure. bioRxiv 2022–07 (2022).

7. Patel, R. et al. Investigating microstructural variation in the human hippocampus using non-negative matrix factorization. Neuroimage 207, 116348 (2020).

8. Maass, A. et al. Laminar activity in the hippocampus and entorhinal cortex related to novelty and episodic encoding. Nat. Commun. 5, 5547, DOI: 10.1038/ncomms6547 (2014).

9. Aitken, F. & Kok, P. Hippocampal representations switch from errors to predictions during acquisition of predictive associations. Nat. Commun. 13, 1–13 (2022).

10. Bussy, A. et al. Hippocampal shape across the healthy lifespan and its relationship with cognition. Neurobiol. Aging 106, 153–168, DOI: https://doi.org/10.1016/j.neurobiolaging.2021.03.018 (2021).

11. Wisse, L. E. et al. Hippocampal subfield volumes at 7t in early alzheimer’s disease and normal aging. Neurobiol. Aging 35, 2039–2045, DOI: https://doi.org/10.1016/j.neurobiolaging.2014.02.021 (2014).

12. Wolf, D., Fischer, F. U., de Flores, R., Chételat, G. & Fellgiebel, A. Differential associations of age with volume and microstructure of hippocampal subfields in healthy older adults. Hum. brain mapping 36, 3819–3831 (2015).

13. Radhakrishnan, H., Stark, S. M. & Stark, C. E. Microstructural alterations in hippocampal subfields mediate age-related memory decline in humans. Front. Aging Neurosci. 12, 94 (2020).

14. Small, S. A., Schobel, S. A., Buxton, R. B., Witter, M. P. & Barnes, C. A. A pathophysiological framework of hippocampal dysfunction in ageing and disease. Nat. Rev. Neurosci. 12, 585–601, DOI: 10.1038/nrn3085 (2011). Number: 10 Publisher: Nature Publishing Group.

15. Blümcke, I. et al. International consensus classification of hippocampal sclerosis in temporal lobe epilepsy: A Task Force report from the ILAE Commission on Diagnostic Methods. Epilepsia 54, 1315–1329, DOI: 10.1111/epi.12220 (2013). _eprint: https://onlinelibrary.wiley.com/doi/pdf/10.1111/epi.12220.

16. Chang, C. et al. The bumps under the hippocampus. Hum. brain mapping 39, 472–490 (2018).

17. Wisse, L. E. et al. A harmonized segmentation protocol for hippocampal and parahippocampal subregions: Why do we need one and what are the key goals? Hippocampus 27, 3–11 (2017).

18. Gross, D. W., Misaghi, E., Steve, T. A., Wilman, A. H. & Beaulieu, C. Curved multiplanar reformatting provides improved visualization of hippocampal anatomy. Hippocampus 30, 156–161 (2020).

19. Duvernoy, H. M., Cattin, F. & Risold, P.-Y. The Human Hippocampus: Functional Anatomy, Vascularization and Serial Sections with MRI (Springer-Verlag, Berlin Heidelberg, 2013), 4 edn.

20. Shing, Y. L. et al. Hippocampal subfield volumes: age, vascular risk, and correlation with associative memory. Front. aging neuroscience 3, 2 (2011).

21. Peters, A., Nawrot, T. S. & Baccarelli, A. A. Hallmarks of environmental insults. Cell 184, 1455–1468, DOI: 10.1016/j.cell.2021.01.043 (2021).

22. Perosa, V. et al. Hippocampal vascular reserve associated with cognitive performance and hippocampal volume. Brain: A J. Neurol. 143, 622–634, DOI: 10.1093/brain/awz383 (2020).

23. Spallazzi, M. et al. Hippocampal vascularization patterns: A high-resolution 7 Tesla time-of-flight magnetic resonance angiography study. NeuroImage. Clin. 21, 101609, DOI: 10.1016/j.nicl.2018.11.019 (2019).

24. Buch, S., Chen, Y., Jella, P., Ge, Y. & Haacke, E. M. Vascular mapping of the human hippocampus using Ferumoxytol-enhanced MRI. NeuroImage 250, 118957, DOI: 10.1016/j.neuroimage.2022.118957 (2022).

25. Detre, J. A., Leigh, J. S., Williams, D. S. & Koretsky, A. P. Perfusion imaging. Magn. resonance medicine 23, 37–45 (1992).

26. Uludağ, K. & Blinder, P. Linking brain vascular physiology to hemodynamic response in ultra-high field mri. Neuroimage 168, 279–295 (2018).

27. Ivanov, D., Poser, B. A., Huber, L., Pfeuffer, J. & Uludağ, K. Optimization of simultaneous multislice epi for concurrent functional perfusion and bold signal measurements at 7t. Magn. resonance medicine 78, 121–129 (2017).

28. Pohmann, R., Speck, O. & Scheffler, K. Signal-to-noise ratio and MR tissue parameters in human brain imaging at 3, 7, and 9.4 tesla using current receive coil arrays. Magn. Reson. Medicine 75, 801–809, DOI: 10.1002/mrm.25677 (2016).

29. DeKraker, J. et al. Automated hippocampal unfolding for morphometry and subfield segmentation with hippunfold. Elife 11, e77945 (2022).

30. DeKraker, J., Lau, J. C., Ferko, K. M., Khan, A. R. & Köhler, S. Hippocampal subfields revealed through unfolding and unsupervised clustering of laminar and morphological features in 3d bigbrain. Neuroimage 206, 116328 (2020).

31. Amunts, K. et al. Bigbrain: an ultrahigh-resolution 3d human brain model. science 340, 1472–1475 (2013).

32. Kashyap, S., Ivanov, D., Havlicek, M., Poser, B. A. & Uludağ, K. Impact of acquisition and analysis strategies on cortical depth-dependent fmri. Neuroimage 168, 332–344 (2018).

33. Andersson, J. L., Skare, S. & Ashburner, J. How to correct susceptibility distortions in spin-echo echo-planar images: application to diffusion tensor imaging. Neuroimage 20, 870–888 (2003).

34. DeKraker, J., Ferko, K. M., Lau, J. C., Köhler, S. & Khan, A. R. Unfolding the hippocampus: An intrinsic coordinate system for subfield segmentations and quantitative mapping. Neuroimage 167, 408–418 (2018).

35. Cembrowski, M. S., Wang, L., Sugino, K., Shields, B. C. & Spruston, N. Hipposeq: a comprehensive rna-seq database of gene expression in hippocampal principal neurons. elife 5, e14997 (2016).

36. Zhao, X. et al. Transcriptional profiling reveals strict boundaries between hippocampal subregions. J. Comp. Neurol. 441, 187–196 (2001).

37. Alkadhi, K. A. Cellular and molecular differences between area ca1 and the dentate gyrus of the hippocampus. Mol. neurobiology 56, 6566–6580 (2019).

38. Podgorny, O. V. & Gulyaeva, N. V. Glucocorticoid-mediated mechanisms of hippocampal damage: Contribution of subgranular neurogenesis. J. Neurochem. 157, 370–392 (2021).

39. Hsu, J. C. et al. Decreased expression and functionality of nmda receptor complexes persist in the ca1, but not in the dentate gyrus after transient cerebral ischemia. J. Cereb. Blood Flow & Metab. 18, 768–775 (1998).

40. Cembrowski, M. S. & Spruston, N. Heterogeneity within classical cell types is the rule: lessons from hippocampal pyramidal neurons. Nat. Rev. Neurosci. 20, 193–204 (2019).

41. Zhang, N., Gordon, M. L. & Goldberg, T. E. Cerebral blood flow measured by arterial spin labeling mri at resting state in normal aging and alzheimer’s disease. Neurosci. & Biobehav. Rev. 72, 168–175 (2017).

42. Guo, X. et al. Asymmetry of cerebral blood flow measured with three-dimensional pseudocontinuous arterial spin-labeling mr imaging in temporal lobe epilepsy with and without mesial temporal sclerosis. J. Magn. Reson. Imaging 42, 1386–1397 (2015).

43. Lieberman, J. et al. Hippocampal dysfunction in the pathophysiology of schizophrenia: a selective review and hypothesis for early detection and intervention. Mol. psychiatry 23, 1764–1772 (2018).

44. Alsop, D. C. et al. Recommended implementation of arterial spin-labeled perfusion mri for clinical applications: A consensus of the ismrm perfusion study group and the european consortium for asl in dementia. Magn. resonance medicine 73, 102–116 (2015).

45. Donahue, M. J., Lu, H., Jones, C. K., Pekar, J. J. & van Zijl, P. C. An account of the discrepancy between mri and pet cerebral blood flow measures. a high-field mri investigation. NMR Biomed. An Int. J. Devoted to Dev. Appl. Magn. Reson. In vivo 19, 1043–1054 (2006).

46. Shaw, K. et al. Neurovascular coupling and oxygenation are decreased in hippocampus compared to neocortex because of microvascular differences. Nat. communications 12, 3190 (2021).

47. Nair, V., Palm, D. & Roth, L. Relative vascularity of certain anatomical areas of the brain and other organs of the rat. Nature 188, 497–498 (1960).

48. Zhang, X. et al. High-resolution mapping of brain vasculature and its impairment in the hippocampus of alzheimer’s disease mice. Natl. Sci. Rev. 6, 1223–1238 (2019).

49. Cavaglia, M. et al. Regional variation in brain capillary density and vascular response to ischemia. Brain research 910, 81–93 (2001).

50. Soltesz, I. & Losonczy, A. Ca1 pyramidal cell diversity enabling parallel information processing in the hippocampus. Nat. neuroscience 21, 484–493 (2018).

51. Michaelis, E. K. Selective neuronal vulnerability in the hippocampus: Relationship to neurological diseases and mechanisms for differential sensitivity of neurons to stress. (2012).

52. Pfeuffer, J. et al. Perfusion-based high-resolution functional imaging in the human brain at 7 tesla. Magn. Reson. Medicine: An Off. J. Int. Soc. for Magn. Reson. Medicine 47, 903–911 (2002).

53. Duvernoy, H. M., Cattin, F. & Risold, P.-Y. The Human Hippocampus: Functional Anatomy, Vascularization and Serial Sections with MRI (Springer Science & Business Media, 2013).

54. Chappell, M. A. et al. Partial volume correction of multiple inversion time arterial spin labeling mri data. Magn. resonance medicine 65, 1173–1183 (2011).

55. Erdem, A., Yaşargil, M. G. & Roth, P. Microsurgical anatomy of the hippocampal arteries. J. neurosurgery 79, 256–265 (1993).

56. Wisse, L. E. et al. Automated hippocampal subfield segmentation at 7t mri. Am. J. Neuroradiol. 37, 1050–1057 (2016).

57. Ivanov, D. et al. Comparison of 3 t and 7 t asl techniques for concurrent functional perfusion and bold studies. Neuroimage 156, 363–376 (2017).

58. Marques, J. P. et al. Mp2rage, a self bias-field corrected sequence for improved segmentation and t1-mapping at high field. Neuroimage 49, 1271–1281 (2010).

59. Eggenschwiler, F., Kober, T., Magill, A. W., Gruetter, R. & Marques, J. P. Sa2rage: A new sequence for fast b1+-mapping. Magn. resonance medicine 67, 1609–1619 (2012).

60. Marques, J. P. & Gruetter, R. New developments and applications of the mp2rage sequence-focusing the contrast and high spatial resolution r1 mapping. PloS one 8, e69294 (2013).

61. Haast, R. A. et al. Effects of mp2rage b1+ sensitivity on inter-site t1 reproducibility and hippocampal morphometry at 7t. Neuroimage 224, 117373 (2021).

62. Chappell, M. A., Groves, A. R., Whitcher, B. & Woolrich, M. W. Variational bayesian inference for a nonlinear forward model. IEEE Transactions on Signal Process. 57, 223–236 (2008).

63. Olsen, R. K. et al. Progress update from the hippocampal subfields group. Alzheimer’s & Dementia: Diagn. Assess. & Dis. Monit. 11, 439–449 (2019).

64. Teeuwisse, W. M., Webb, A. G. & van Osch, M. J. Arterial spin labeling at ultra-high field: all that glitters is not gold. Int. J. Imaging Syst. Technol. 20, 62–70 (2010).

65. Teeuwisse, W. M., Brink, W. M. & Webb, A. G. Quantitative assessment of the effects of high-permittivity pads in 7 tesla mri of the brain. Magn. resonance medicine 67, 1285–1293 (2012).

66. Kashyap, S. et al. The impact of b1+ on the optimisation of high-resolution asl acquisitions at 7t. In Proc. Intl. Soc. Mag. Reson. Med., vol. 29, 1417 (2021).

67. Hurley, A. C. et al. Tailored rf pulse for magnetization inversion at ultrahigh field. Magn. Reson. Medicine: An Off. J. Int. Soc. for Magn. Reson. Medicine 63, 51–58 (2010).

68. Petr, J. et al. Effects of systematic partial volume errors on the estimation of gray matter cerebral blood flow with arterial spin labeling mri. Magn. Reson. Mater. Physics, Biol. Medicine 31, 725–734, DOI: 10.1007/s10334-018-0691-y (2018).

69. Günther, M., Oshio, K. & Feinberg, D. A. Single-shot 3d imaging techniques improve arterial spin labeling perfusion measurements. Magn. Reson. Medicine 54, 491–498, DOI: 10.1002/mrm.20563 (2005).

70. Vidorreta, M. et al. Evaluation of segmented 3d acquisition schemes for whole-brain high-resolution arterial spin labeling at 3t. NMR Biomed. 27, 1387–1396, DOI: https://doi.org/10.1002/nbm.3201 (2014).

71. Kurban, D. et al. High-resolution perfusion and blood-volume fmri at 7t with simultaneous multi-slice spiralout acquisitions. In Proceedings of the 37th Annual Scientific Meeting, ESMRMB Online (2020).

72. Kashyap, S. et al. Sub-millimetre resolution laminar fmri using arterial spin labelling in humans at 7 t. Plos one 16, e0250504 (2021).

73. Woods, J. G., Chappell, M. A. & Okell, T. W. A general framework for optimizing arterial spin labeling mri experiments. Magn. Reson. Medicine 81, 2474–2488, DOI: 10.1002/mrm.27668 (2019).

74. Fan, A. P. et al. Long-delay arterial spin labeling provides more accurate cerebral blood flow measurements in moyamoya patients. Stroke 48, 2441–2449, DOI: 10.1161/STROKEAHA.117.017773 (2017). https://www.ahajournals.org/doi/pdf/10.1161/STROKEAHA.117.017773.

75. Johnson, A. C. Hippocampal vascular supply and its role in vascular cognitive impairment. Stroke 54, 673–685 (2023).

76. Haines, K., Smith, N. & Webb, A. New high dielectric constant materials for tailoring the b1+ distribution at high magnetic fields. J. magnetic resonance 203, 323–327 (2010).

77. Haast, R. A., Ivanov, D. & Uludağ, K. The impact of correction on mp2rage cortical t 1 and apparent cortical thickness at 7 t. Hum. brain mapping 39, 2412–2425 (2018).

78. Hennig, J., Nauerth, A. & Friedburg, H. Rare imaging: a fast imaging method for clinical mr. Magn. resonance medicine 3, 823–833 (1986).

79. Axel, L. Blood flow effects in magnetic resonance imaging. Magn. resonance annual 237–244 (1986).

80. Parker, D. L., Yuan, C. & Blatter, D. D. Mr angiography by multiple thin slab 3d acquisition. Magn. resonance medicine 17, 434–451 (1991).

81. Wehrli, F. W. Time-of-flight effects in mr imaging of flow. Magn. resonance medicine 14, 187–193 (1990).

82. Kwong, K. K. et al. Mr perfusion studies with t1-weighted echo planar imaging. Magn. Reson. Medicine 34, 878–887 (1995).

83. Kim, S.-G. Quantification of relative cerebral blood flow change by flow-sensitive alternating inversion recovery (fair) technique: application to functional mapping. Magn. resonance medicine 34, 293–301 (1995).

84. Wong, E. C., Buxton, R. B. & Frank, L. R. Quantitative imaging of perfusion using a single subtraction (quipss and quipss ii). Magn. resonance medicine 39, 702–708 (1998).

85. Shaw, T. B. et al. Non-linear realignment improves hippocampus subfield segmentation reliability. NeuroImage 203, 116206 (2019).

86. Kashyap, S. srikash/presurfer: ondu, DOI: 10.5281/zenodo.4626841 (2021).

87. Dale, A. M., Fischl, B. & Sereno, M. I. Cortical surface-based analysis: I. segmentation and surface reconstruction. Neuroimage 9, 179–194 (1999).

88. Aguirre, G. K., Detre, J. A., Zarahn, E. & Alsop, D. C. Experimental design and the relative sensitivity of bold and perfusion fmri. Neuroimage 15, 488–500 (2002).

89. Liu, T. T. & Wong, E. C. A signal processing model for arterial spin labeling functional mri. Neuroimage 24, 207–215 (2005).

90. Rane, S. D. & Gore, J. C. Measurement of t1 of human arterial and venous blood at 7 t. Magn. resonance imaging 31, 477–479 (2013).

91. Panchuelo, R. M. S., Mougin, O., Turner, R. & Francis, S. T. Quantitative t1 mapping using multi-slice multi-shot inversion recovery epi. NeuroImage 234, 117976 (2021).

92. Yushkevich, P. A., et al. Fast automatic segmentation of hippocampal subfields and medial temporal lobe subregions in 3 tesla and 7 tesla t2-weighted mri. Alzheimer’s & Dementia 7, P126–P127 (2016).

93. Greve, D. N. & Fischl, B. Accurate and robust brain image alignment using boundary-based registration. Neuroimage 48, 63–72 (2009).

94. Yushkevich, P. A. et al. User-guided 3D active contour segmentation of anatomical structures: Significantly improved efficiency and reliability. Neuroimage 31, 1116–1128 (2006).

95. Köster, J. & Rahmann, S. Snakemake—a scalable bioinformatics workflow engine. Bioinformatics 28, 2520–2522 (2012).

96. Marcus, D. et al. Informatics and data mining tools and strategies for the human connectome project. Front. neuroinformatics 5, 4 (2011).

97. Gerig, G., Kubler, O., Kikinis, R. & Jolesz, F. A. Nonlinear anisotropic filtering of mri data. IEEE Transactions on medical imaging 11, 221–232 (1992).

98. Weickert, J. Coherence-enhancing diffusion filtering. Int. journal computer vision 31, 111–127 (1999).

99. Gulban, O. F., Schneider, M., Marquardt, I., Haast, R. A. & De Martino, F. A scalable method to improve gray matter segmentation at ultra high field mri. PloS one 13, e0198335 (2018).

100. Ritter, F. et al. Medical image analysis. IEEE pulse 2, 60–70 (2011).

101. Selle, D., Preim, B., Schenk, A. & Peitgen, H.-O. Analysis of vasculature for liver surgical planning. IEEE transactions on medical imaging 21, 1344–1357 (2002).

102. Mattern, H. Book of abstracts ESMRMB 2021 online 38th annual scientific meeting 7-9 october 2021. MAGMA 34, 1–204 (2021).

103. Brett, M. et al. nipy/nibabel: 2.4.1, DOI: 10.5281/zenodo.3233118 (2019).

104. Hagberg, A., Swart, P. & S Chult, D. Exploring network structure, dynamics, and function using networkx. Tech. Rep., Los Alamos National Lab.(LANL), Los Alamos, NM (United States) (2008).

105. Vallat, R. Pingouin: statistics in python. J. Open Source Softw. 3, 1026, DOI: 10.21105/joss.01026 (2018).

106. Markello, R. D. & Misic, B. Comparing spatial null models for brain maps. NeuroImage 236, 118052 (2021).

107. Virtanen, P. et al. SciPy 1.0: Fundamental Algorithms for Scientific Computing in Python. Nat. Methods 17, 261–272, DOI: 10.1038/s41592-019-0686-2 (2020).

